# Quorum-Sensing-Mediated Extracellular Electron Transfer Enables Hydrogel Morphogenesis

**DOI:** 10.64898/2026.06.16.732721

**Authors:** Ismar E. Miniel Mahfoud, Vidhika S. Damani, Gina Partipilo, Andy Y. Liu, Benjamin K. Keitz

## Abstract

Engineered living materials seek to capture the sensitivity, responsiveness, and programmable characteristics of biological systems. One emergent property of living systems is genetically driven spatial patterning, which controls cell differentiation and the development of complex multicellular organisms. Inspired by this capability, we use bacteria to spatially control material assembly. In our system, extracellular electron transfer (EET) flux from *Shewanella oneidensis* drives hydrogel synthesis via copper-catalyzed radical polymerization. We first construct a recombinant quorum sensing system in *S. oneidensis* that allows for cell-cell communication between “sender” and “receiver” cells through an autoinducer. We then examine controlled gene expression and EET-driven chemical transformation in various synthetic consortia. Via diffusion through agarose, we examine 2D patterns of gene expression relative to localized sender cell populations and demonstrate controlled hydrogel crosslinking in predictable patterns. Finally, we apply computational methods and NOT logic in “receiver” cells towards more complex patterns of gene expression. Our results highlight the potential of bacteria to program material systems with life-like properties including self-assembly, environmental responsiveness, and patterned differentiation.

## Introduction

Across the tree of life, patterns including symmetry^1^, stripes^2,3^, tessellations^4^, and spirals^5^ have arisen as a result of spatiotemporal activation and inhibition of biochemical signals^6^. The genetically encoded formation of these patterns allows for the emergence of self-assembly^7^, metabolic regulation^8^, self-healing^9^, and multicellularity^10–12^. Understanding the underlying genetic, metabolic, and regulatory mechanisms behind pattern formation and population dynamics is also useful in combating phenomena like congenital anomalies^13,14^. Most patterns in living systems are formed through the genetic interpretation of morphogen signals, as proposed in Wolpert’s “French flag” model^15,16^. In short, cells produce diffusible signals that surrounding cells genetically interpret using the concentration gradient of the signal to create distinct spatiotemporal phenotypes^2,6,15,17–19^. Similar patterning behavior can be recapitulated in bacteria using quorum sensing. For example, quorum sensing bacteria continuously produce, sense, and genetically respond to autoinducer molecules^20–22^. As population size increases, the concentration of the autoinducer in the immediate environment passes a threshold, leading to the expression of key genes related to functions such as virulence and biofilm formation by the community^23–27^. The underlying principle connecting quorum sensing and morphogen systems lies in the secretion of a diffusible signal and a concentration-dependent phenotypical response. Bacterial systems are especially useful in studying and predicting patterning phenomena due to advances in synthetic biology and genetic logic that allow for tunable genetic responses to signals^28–30^. Consequently, the pairing of multiple, orthogonal quorum sensing signals and synthetic gene circuits has resulted in numerous, sophisticated examples of patterned gene expression in bacterial populations^31–40^. However, the vast majority of these examples have focused on patterning the expression of fluorescent proteins and there are relatively few examples of translating spatial differences in bacterial protein expression into material properties, such as stiffness or color^41^.

The growing field of engineered living materials (ELMs) seeks to co-opt the properties of self-organization, patterning, and other properties of living systems for the development of next-generation synthetic, semisynthetic, and biological materials. Some ELMs use living cells as sensors and actuators, allowing for growth and adaption while performing tailored functions beyond the cells’ native biological abilities^42–44^. One particularly useful class of materials to endow with such properties is hydrogels, which are liquid-swollen polymer networks with extensive applications in tissue engineering^45–49^, 3D printing^50,51^, drug delivery^45,46,52,53^, biosensing^54,55^, and soft robotics^56–58^. The porous, aqueous structure of hydrogels is ideal for embedding microorganisms that are genetically tractable and sensitive to their environment. For example, hydrogels containing yeast or bacteria can undergo biomass-dependent shape changes while remaining genetically tractable and responsive for applications such as soft actuation^59^, bacterial capture or containment^60^, and cargo delivery^61,62^. Some bacteria can generate their own matrices through biofilm formation or curli fiber secretion, but these natural materials can be limited in functionality, robustness, and scalability^63–67^. Programming the mechanical characteristics of hydrogels and other materials using synthetic biology is an attractive possibility for living materials, but is limited by a lack of mechanisms that connect gene expression to material properties.

Here, we use synthetic gene circuits relying on quorum sensing signals to spatiotemporally control hydrogel mechanical properties. By controlling the expression of key EET proteins using genetic circuitry, we developed hydrogels whose stiffness is determined by bacterial genetic responses to one or more chemical inputs^73,74^. Leveraging this capability, we first develop a tunable recombinant quorum sensing system in *Shewanella oneidensis* that recognizes quorum sensing molecules and regulates the expression of *sfgfp* (**Fig. 1**). We extend this regulation to key EET genes and achieve transcriptional control over EET-driven biocatalytic and material crosslinking applications. To further increase regulation capacity, we introduce an additional regulatory layer by placing the autoinducer synthase under inducible control. Using immobilized “sender” and “receiver” cells, we control pattern formation relative to the source of the sender signal and demonstrate that the patterned expression of EET proteins leads to spatial control EET-driven material changes. Based on experimental data, we developed a computational tool for *in silico* design of patterns that can then be realized as fields of biocatalytic electroactivity. Finally, we explored methods of creating more complex gene expression pattens by constructing NOT logic gates and combining receiver strains in co-culture. Overall, we demonstrate a system for tunable and predictable patterned material formation as a result of biotic interpretation of spatiotemporal environmental signals, with potential applications in 3D bioprinting, tissue engineering, environmental sensing, and modeling developmental events.

**Figure 1.**
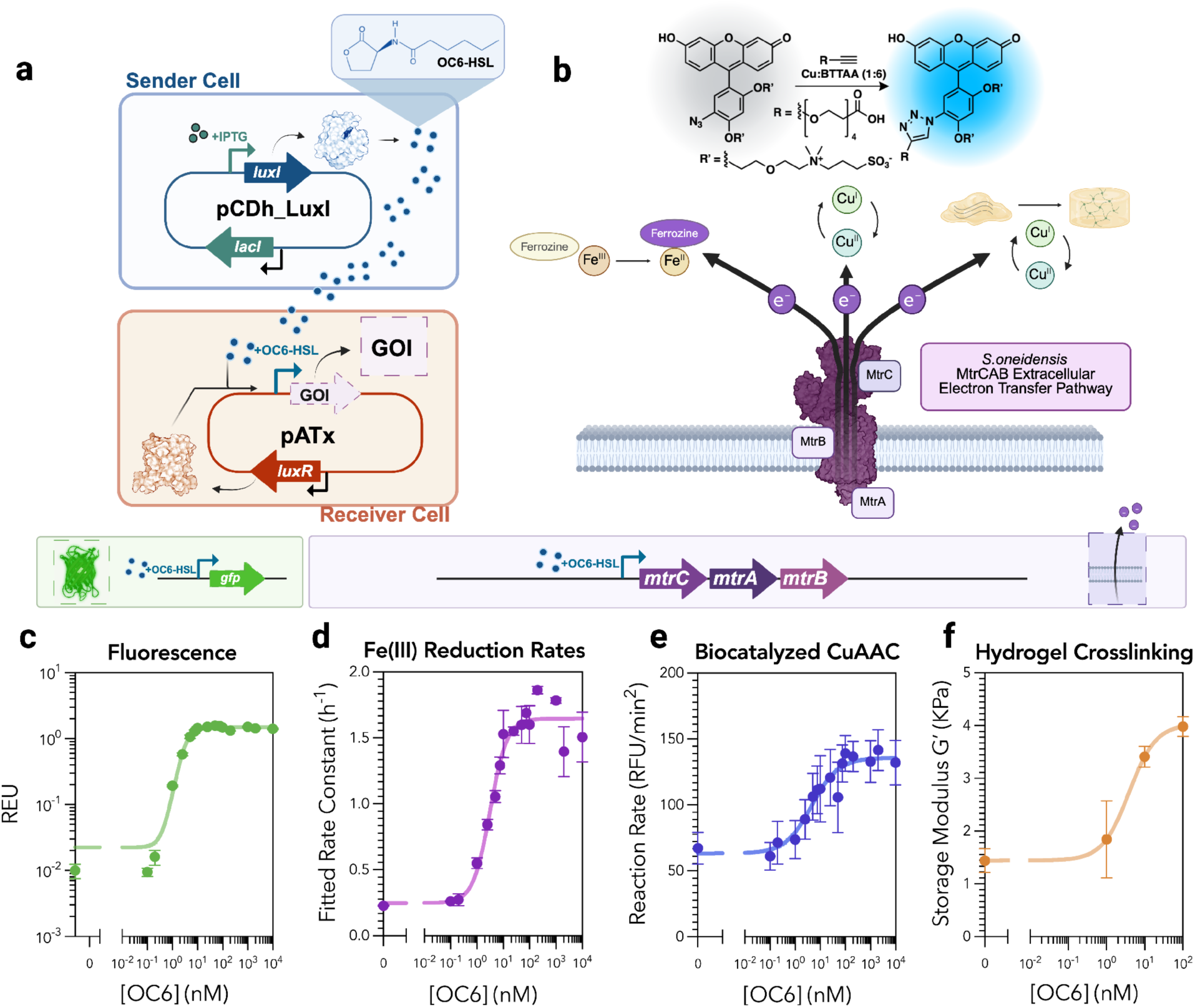
Construction and characterization of recombinant quorum sensing in *S.oneidensis*. **a.** Schematic illustrating the plasmid-based expression of the canonical LuxIR quorum sensing system in wild-type *S. oneidensis*. “Sender” cells contain the AHL-synthase gene LuxI under control of the IPTG-inducible P_tac_ promoter. LuxI generates OC6 that diffuses out of the cell. “Receiver” cells recognize OC6 and activate transcription of a GOI downstream of the pLux promoter **b.** Schematic illustrating extracellular electron transfer through the MtrCAB protein pathway in *S.oneidensis*. EET flux can be applied for colorimentric measurements of Fe(III) reduction, CuAAC chemistry, and radical polymerization. **c.** REU response function of *S.oneidensis* MR-1 expressing the OC6-inducible pAT3 *sfgfp* buffer gate. **d.** Anaerobic Fe(III) reduction rate response function of aerobically pregrown *S.oneidensis ΔMtr* expressing the OC6-inducible *mtrCAB* buffer gate (pAT4). **e.** Rate of CuAAC between 0.6 μM CalFluor 488 and 100 μM alkyne-PEG4-acid and 50 μM Cu:BTTAA (1:6) in minimal media as a function of variably induced *S. oneidensis ΔMtr* containing pAT4. **f.** Storage moduli of 5 wt% methacrylated PEG and 1 wt% acrylated agarose co-polymer networks cross-linked via variably induced *S.oneidensis ΔMtr* containing pAT4 driving atom-transfer radical polymerization (ATRP). Moduli were measured using shear rheometry after swelling. For each graph individual replicates were fit to a four-parameter activating Hill function with weighted error. All data represent the mean ± SEM of the appropriate measurement for n = 3 biological replicates. Created using Biorender.com

## Results

### Recombinant Quorum Sensing in *S. oneidensis*

Towards achieving patterned gene expression of proteins critical for EET, we developed recombinant quorum sensing in *S.oneidensis*^75,76^. We chose *S.oneidensis* and transcriptional EET regulation because this species is genetically tractable and controlled *mtrCAB* expression directly drives EET rates. Our system relies on the well-characterized LuxIR system from *Allivibrio fischerii*, where it natively controls fluorescence dependent on the local concentration of autoinducer produced by the population. We first attempted single plasmid self-patterning approaches^77,78^ but found they were not as stable or robust in *Shewanella* for the purposes of our system. Inspired by more complex patterning systems^31^ we decided on a “sender” and “receiver” cell set-up. As depicted in **Fig. 1a**, “sender” cells contain a plasmid on which the LuxI AHL synthase gene is under the control of the LacI-*P_tacsymO_* regulator-promoter pair. In the presence of isopropyl-β-thiogalactopyranoside (IPTG), LuxI is expressed, and synthesizes a six-carbon chain homoserine lactone (OC6-HSL). Following biosynthesis, OC6 diffuses out of the sender cells and serves as an inducer to the “receiver” cells.

Our suite of “receiver” *S. oneidensis* cells contain the pATx plasmid backbone, where the transcriptional activator LuxR is constitutively expressed. Upon binding to OC6, LuxR dimerizes and activates transcription of genes downstream of its cognate *P_lux_* promoter^79^. We began by parameterizing the behavior of receiver cells using wild-type *S*. *oneidensis* MR-1 containing the pAT3 plasmid, which expresses *sfgfp* as a reporter of *P_lux_* promoter activation (**Fig. 1b**). Fluorescence was measured after overnight growth in minimal media containing variable amounts of OC6 and was quantified via relative expression units (REUs), which aid in comparing the performance of transcriptional circuits regulating RNA polymerase activity^30^. REU values were determined by normalizing measurements to fluorescence from an *S. oneidensis* strain carrying a plasmid with constitutive *sfgfp* expression driven by the *P_trc*_* promoter.

The fluorescent pAT3 plasmid confirmed that the regulatory gene architecture of the pATx plasmid backbone yields the expected concentration-dependent behavior resulting in an activating Hill Function (**Fig. 1c**). The same backbone was used to construct and transform the pAT4 plasmid into *S. oneidensis* JG1194 which has genomic knockouts of the Mtr pathway and various accessory proteins that facilitate EET. The Mtr pathway is the main molecular mechanism of EET used by *Shewanella* and it consists of membrane-bound cytochromes MtrA and MtrC which span the outer membrane via the beta-barrel protein MtrB. The pAT4 plasmid places polycistronic expression of genes *mtrC*, *mtrA,* and *mtrB* under P*_lux_* control (**Fig. 1d-f**). To initially measure EET rates relative to pAT4 induction, we used a previously established in-situ Fe(III)-reduction assay^74^. Here, EET-driven Fe(III) reduction is tracked colorometrically using ferrozine, a color-shifting Fe(II) chelator. The rate of iron reduction at each induction level was plotted against the concentration of OC6 and revealed a characteristic activating Hill function, confirming MtrCAB expression was under transcriptional control (**Fig. 1d**).

To highlight the tunable biocatalytic applications of EET using the pATx backbone, we next performed a similar assay where the terminal electron acceptor was a Cu-BTTAA catalyst-ligand complex. This complex, when reduced by EET, drives copper-catalyzed alkene-azide cycloaddition (CuAAC)^80^. In this case, successful CuAAC resulted in the formation of a fluorescent chemical probe. After tracking the fluorescence and calculating rates of conversion using a second-order polynomial fit, reaction rates were plotted against the OC6 concentration. Similar to our previous observation, we measured the expected activating Hill function behavior (**Fig. 1e**).

After validating the genetic architecture, heterologous EET-pathway expression, and biocatalytic ability of our engineered strain, we measured its control over a material system. To do so, we used an acrylated agarose (ACAG) (**Fig. S12)** and methacrylated 4-arm PEG (PEGMA) system where chemical crosslinking is controlled by copper-catalyzed radical polymerization. EET flux to the copper catalyst controls the polymerization rate and overall mechanical properties of the hydrogel via an atom-transfer radical polymerization (ATRP) mechanism^81^. After a 2-hour anaerobic incubation with induced bacteria, the hydrogels were swollen in PBS overnight and their storage modulus was determined using a frequency sweep on a shear rheometer. The stiffness of the gels shows the expected Hill function behavior relative to OC6 levels for 5% PEGMA and 1% ACAG formulations (**Fig. 1f**). A hydrogel composed of only PEGMA showed the same behavior (**Fig. S1**). Together, these results demonstrate the OC6-dependent gene expression of multiple genes of interest in *Shewanella*, and the ability of tunable EET expression to control a variety of orthogonal chemistries.

### Cocultured Patterning Strains Allow for Tunable Control of Various Genetic Outputs and Chemistries

Population size and biologically produced autoinducer concentrations are key variables in determining genetic outputs of quorum sensing populations. After validating the receiver cell response to exogenous OC6 using lab stocks, we validated our strains’ behavior as a quorum sensing system in defined liquid co-cultures. We examined the effect population and gene expression on the final genetic output of our synthetic consortia by combinatorially varying the initial sender:receiver population ratio and controlling LuxI expression using IPTG (**Fig. 2a**). We first tested the pAT3 fluorescent receivers, and recorded biomass-normalized fluorescence values from each co-culture condition after overnight incubation (**Fig. 2b, S2**). As expected, higher IPTG concentrations led to more LuxI expression by sender cells, thus increasing the OC6 concentration and the amount of *sfgfp* expressed by pAT3 receiver cells. The ratio of sender to receiver cells also played a role in the final response; a higher ratio of sender:receiver cells produced higher fluorescence values, even in conditions with minimal to no IPTG. This observation can be explained by the sensitivity of the LuxIR system, where background levels of leaky expression of LuxI can result in a detectable signal, especially in the small, high cell density volumes used in our high-throughput 96-well plate characterization. Visualized as a heatmap (**Fig. 2b)**, the trend of increasing output as a result of more sender cells and higher IPTG concentrations validate the function and tunability of our engineered quorum-sensing system.

**Figure 2.**
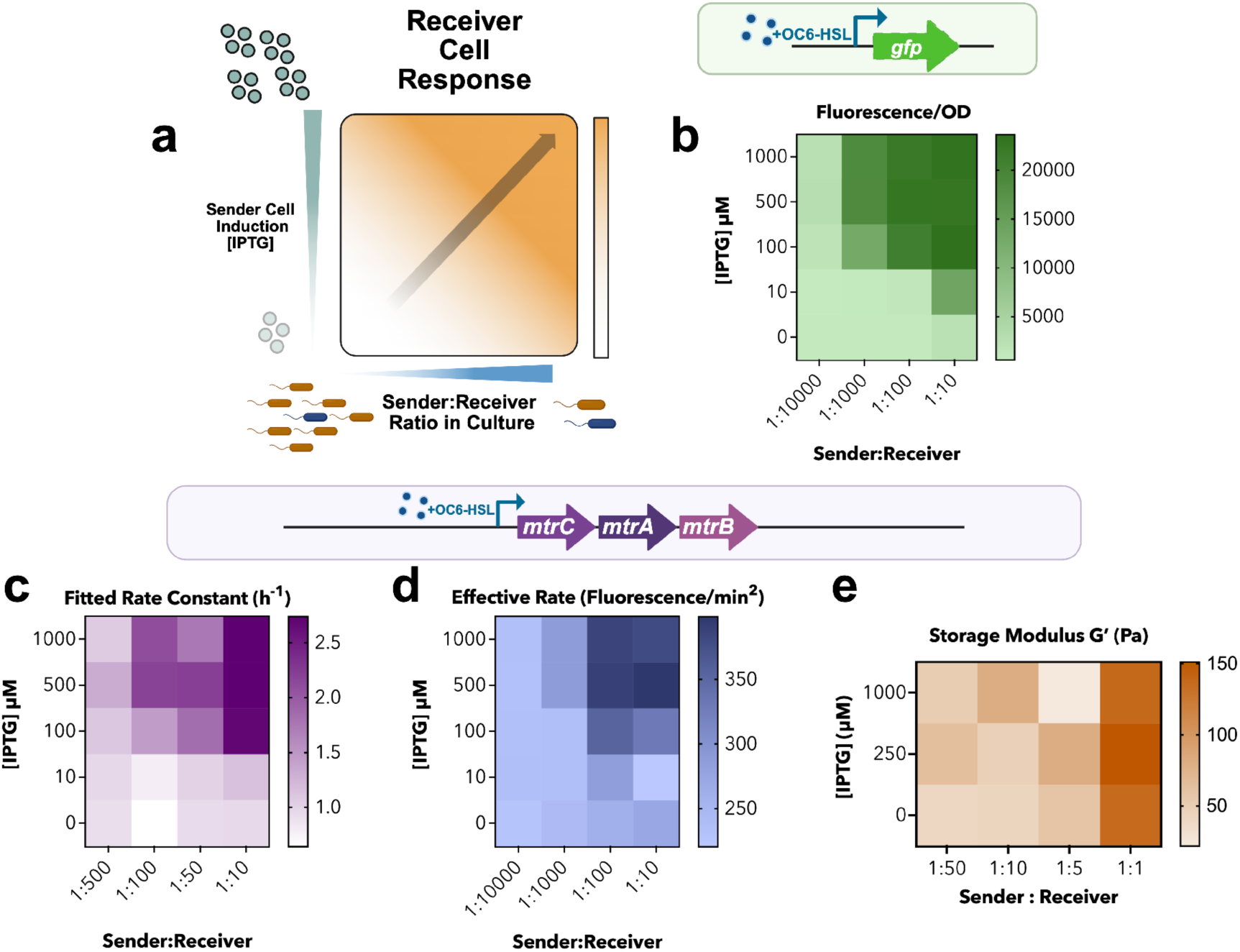
Cocultured patterning strains allow for tunable control of various genetic outputs and chemistries. **a.** Schematic illustrating the experimental parameters of sender and receiver cell cocultures used to examine the tunable transcriptional space of our recombinant quorum sensing Shewanella communities. **b.** Endpoint fluorescence/OD600 response of sender and receiver cocultures of *S. oneidensis* MR-1 containing the pCDh_LuxI and pAT3 plasmids, respectively. **c.** Fe(III) reduction rates of cocultures of sender and receiver *S. oneidensis ΔMtr* containing the pCDh_LuxI and pAT4 plasmids, respectively. **d.** CuAAC biocatalysis rates of cocultures of sender and receiver *S. oneidensis ΔMtr* containing the pCDh_LuxI and pAT4 plasmids, respectively. **e.** Storage modulus of 5 wt% PEGMA polymer networks crosslinked via cocultures of sender and receiver *S. oneidensis ΔmtrCAB* containing the pCDh_LuxI and pAT4 plasmids, Data for **b-d** represent the mean of the appropriate measurement for n = 3 biological replicates, 1 biological replicate for **e.** Created using Biorender.com

Having confirmed that this assay can capture the quorum-sensing behavior of our synthetic system, we repeated this assay with receiver cells expressing MtrCAB. To ensure any observed EET activity was a result of cell-cell communication through OC6-HSL and not native EET processes, the sender cell plasmid was transformed into a ΔMtr background strain. Instead of measuring fluorescence, we used the sender:receiver mixtures to seed a ferrozine/Fe(III) mixture containing varying amounts of IPTG. We measured the iron reduction rate of each condition and observed the same trends as the fluorescence protocol, mainly iron reduction increased with the concentration of IPTG (**Fig. 2c, S3).** We then applied transcriptional regulation through community dynamics by using the sender:receiver mixtures to catalyze CuAAC reactions containing varying concentrations of IPTG. The co-culture of sender and receiver cells successfully catalyzed the formation of product and the plotted conversion rates (determined via a fluorescence assay) also showed the expected trend (**Fig. 2e**). Finally, we performed crosslinking of 5 wt% PEGMA with variably induced sender and receiver co-cultures. Generally, higher sender cell induction and abundance resulted in the stiffest moduli (∼150 Pa), consistent with the highest moduli achieved with exogenous OC6 in previous experiments (**Fig. S1).** These results demonstrate transcriptional regulation through community dynamics for various genetic and chemical outputs using recombinant quorum sensing in *S.oneidensis*.

### Spatiotemporal Expression Patterns are Consistent Across Genetic Outputs and Chemistries

Having validated controlled cell-cell communication between sender and receiver cells, we explored whether gene expression could be translated to two-dimensional pattern formation. These assays were performed on 60 mm plates of agarose in minimal media. The receiver cell lawn was created by plating low weight percent agar seeded with receiver cells, and induced sender cells were localized onto a filter paper disc. As the plates incubate, the sender cells produce OC6 that diffuses radially on the receiver cell lawn. The growing receiver cells then express the GOI in response to the OC6 gradient. The size of the radial pattern is determined by the variable concentration of IPTG in the sender cell population, which ultimately controls the amount of OC6 produced (**Fig. 3a**).

**Figure 3.**
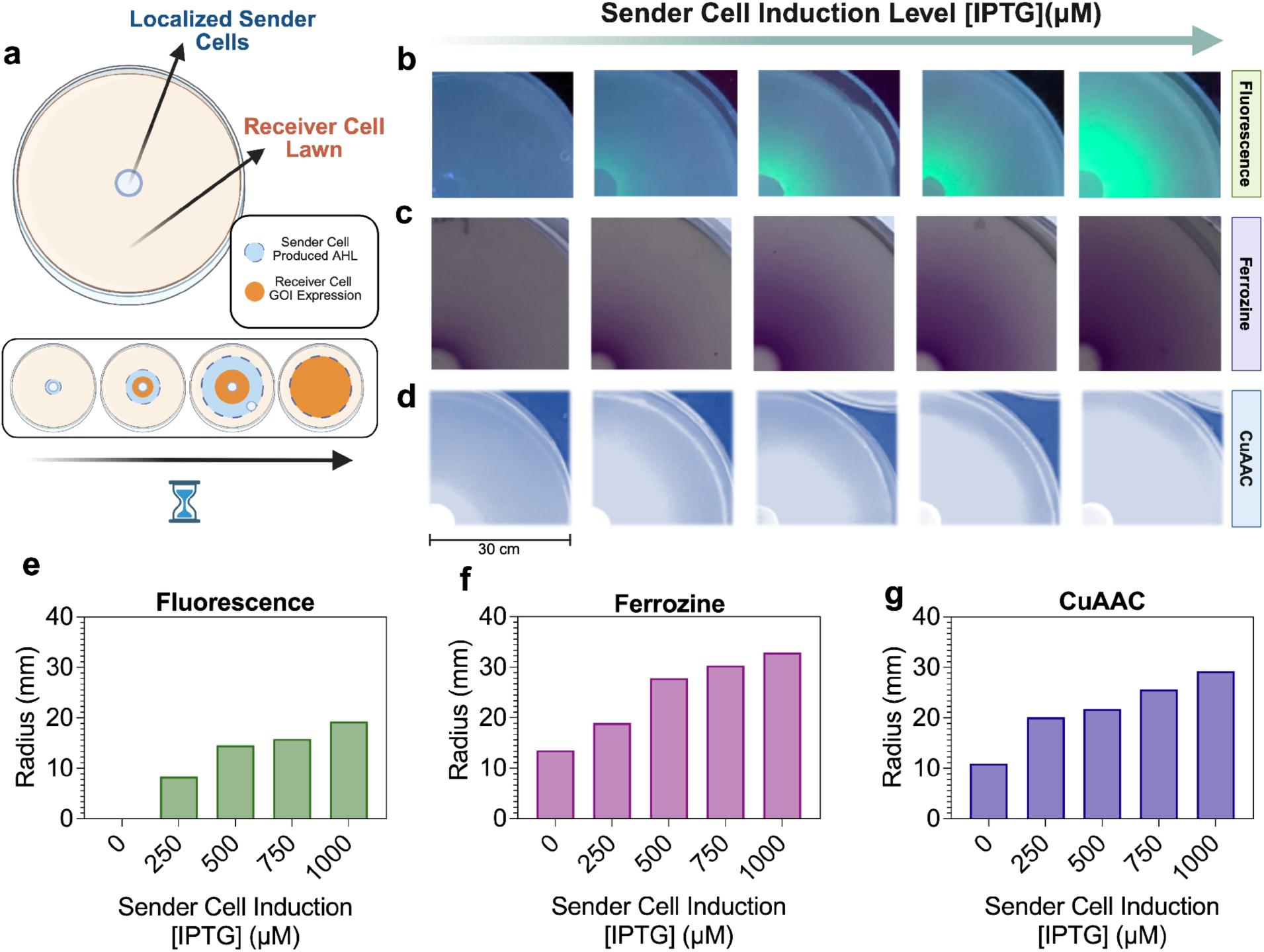
Spatiotemporal expression patterns are consistent across genetic outputs and chemistries. **a.** Schematic depicting the experimental setup for two-dimensional pattern formation on a 60 mm minimal media agarose plate with a 0.7% top agar lawn of receiver cells. Sender cells concentrated and localized on filter paper with higher IPTG concentrations create larger patterns of expression in the receiver cell lawn. **b.** Patterns of GFP expression as visualized on an UV transilluminator following 18 hour incubation at 30°C. **c.** Patterns of MtrCAB expression as visualized through a ferrozine pourover. 2mM Fe(II) citrate and 1 mM ferrozine in .7% agarose was plated on top of the receiver lawn after 18 hours of incubation at 30°C and imaged after 10 minutes on a white light box. **d.** Patterns of MtrCAB expression applied to patterned biocatalyzed CuAAC chemistry conversion of precursors into a fluorescent probe. 0.7% agar in minimal media containing CuAAC reaction components was poured over the receiver lawn after 18 hour incubation at 30°C and was allowed to incubate anaerobically at 30°C for 1 hour. **e-g.** Image analysis of pattern radii corresponding to sender cell induction. Scale bars are 30 mm. Circles seen in the middle of the GFP and ferrozine patterns are a result of removing the sender cell saturated filter paper from the soft lawn agar. Scale bar = 30 mm.

We first tested patterns of GFP expression using the pAT3 receiver cells. After aerobic incubation, fluorescence was captured using an UV transilluminator. As expected, higher levels of sender cell induction resulted in higher fluorescence further from the sender cells (**Fig. 3b, S5**). The radial diffusion of the OC6 signal coming from the sender cells is evident from the tapering off of fluorescence at the outer edge of the pattern due to the gradual decrease in local inducer concentration.

Next, we examined whether we could pattern EET activity in a lawn of pAT4 receiver cells by controlling where the MtrCAB pathway is expressed. The experiment was set up as previously described, but using LuxI sender cells in a ΔMtr strain to avoid any background electroactivity outside of the receiver cell population. After aerobic incubation, the pattern of EET activity was visualized using a modified version of the ferrozine assays (previously used to measure iron reduction rates). By pouring low weight percent agarose containing Fe(III) citrate and ferrozine in a 2:1 molar ratio, we visualized where EET is occurring via the purple color change characteristic of Fe(II) reduction **(Fig. S6).** After 10 minutes, distinct patterns of iron reduction and thus EET activity due to MtrCAB expression was observed. The pattern size was consistent with the fluorescence characterization and increased with the IPTG concentration of the sender cells (**Fig. 3c, S7**).

Finally, to determine the functionality of the patterned EET agarose, we used the biocatalyzed CuAAC reaction as an application of “transferring” gene expression patterns to materials and other chemical systems. Here, the CuAAC reaction reagents were added into low-percent minimal media agar, poured on and incubated at 30 °C. The resulting CuAAC-formed fluorescent probe was visualized using a UV transilluminator **(Fig. S8).** Excitingly, biocatalysis only occurred where EET expression was patterned with the same trends observed in the fluorescence and ferrozine assays (**Fig. 3d, S9**). Together, these observations demonstrate tunable control of patterned gene expression in *S.oneidensis* hydrogels with programmed regions of electroactivity used to drive biocatalyzed reactions in specific spatial configurations.

### Hydrogel Crosslinking Patterned by Cell-Cell Communication in *S.oneidensis*

Next, we adapted the two-dimensional sender receiver system for patterned biocatalysis of living hydrogels. These experiments were performed with 1 wt% acrylated agarose and 5 wt% PEGMA hydrogels (**Fig. 4a)**. We first measured patterned crosslinking performed by receiver cells relative to a sender cell population. For these experiments, the hydrogel precursor was seeded with pAT4 receiver cells and cast as strips in a PDMS mold, after which variably induced sender cell populations were placed on the edge of the hydrogel strips. The mold was placed between two glass slides, wrapped in parafilm, and allowed to incubate anaerobically for 18 hours in a humidified chamber (**Fig. 4b**). For rheological measurements, 8 mm discs were punched out of the gel strip and allowed to equilibrate overnight in PBS. Storage modulus (G’) measurements as a function of distance from the sender populations revealed patterns of stiffness and spatial organization dependent on sender cell induction levels. The 250 µM IPTG gel condition had the clearest decreasing trend over the length of the gel robust gradient formation. In contrast, the 1000µM IPTG gel condition had the highest overall stiffness, but variation between excised samples indicate weak spatial organization (**Fig. 4c).**

**Figure 4.**
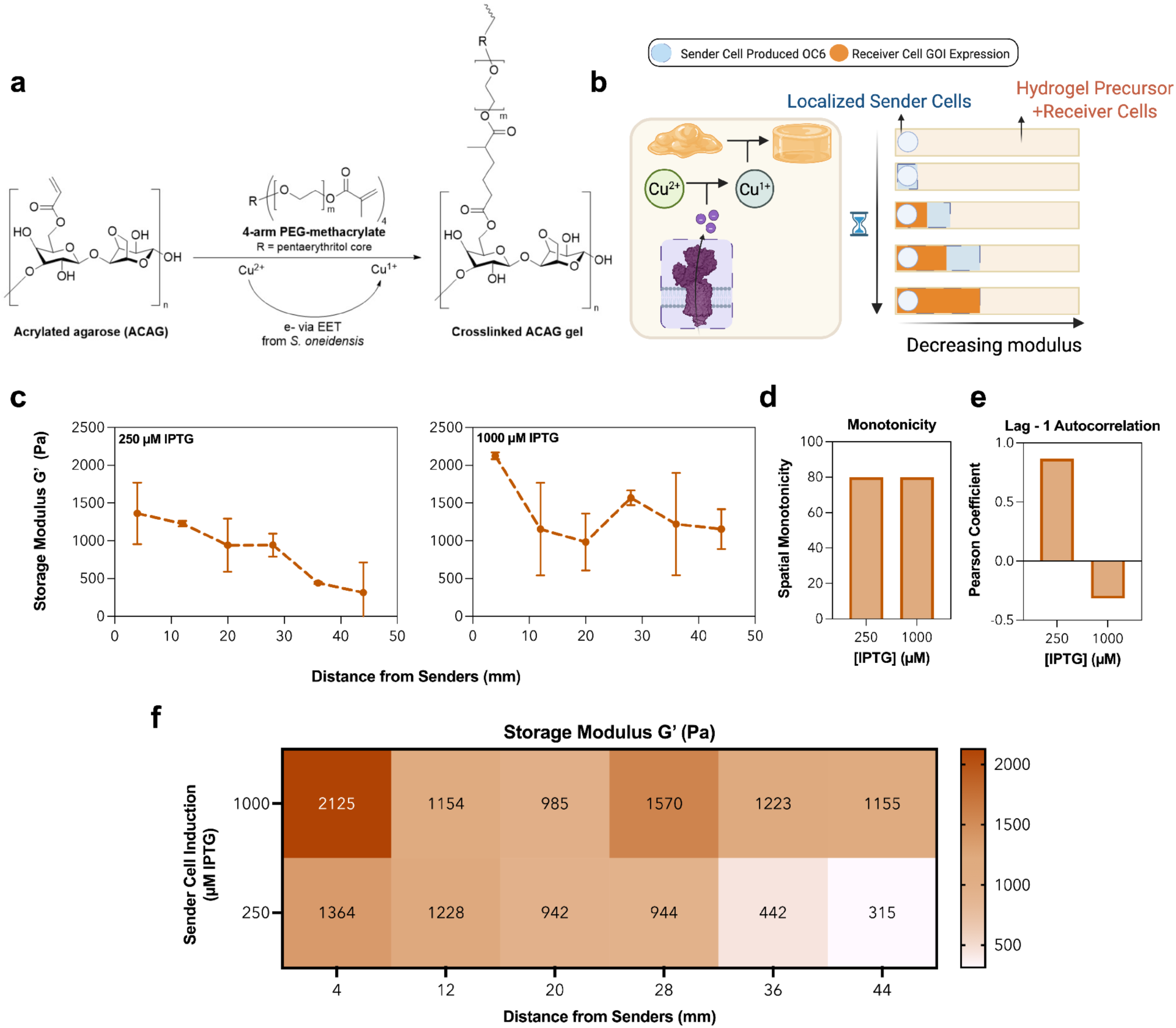
Hydrogel crosslinking patterned by cell-cell communication in *S.oneidensis*. **a.** Schematic of ACAG/PEGMA hydrogel crosslinking through copper-catalyzed ATRP**. b.** Schematic depicting the expected mechanism of pattern formation by lateral diffusion of OC6 in the ACAG/PEGMA hydrogel precursor. **c.** Storage modulus of 1 wt% ACAG and 5 wt% PEGMA hydrogels as a function of distance from the sender cell population **d.** Monotonicity score of hydrogel stiffness as a measure of consistency or smoothness of the observed trend across the entire hydrogel strip**. e.** Lag -1 autocorrelation analysis of hydrogel stiffness as distance from the sender cell population increases. The Pearson coefficient quantifies if the change in modulus across the hydrogel strip follows the expected (decreasing) trend**. f.** Heatmap summarizing both conditions. *n* = 2 biological replicates for patterned hydrogel strips.

The differences between conditions are further quantified using monotonicity scores and lag-1 autocorrelation analysis (**Fig. 4d,e)**. Monotonicity was calculated by giving a score of 1 if adjacent samples from left to right show a decrease in value and a value 0 if not. The score is then divided by the number of adjacent samples to give the monotonicity percentage. In short, it quantifies if the hydrogel overall is decreasing in stiffness as distance from senders increases. Both 250 µM and 1000 µM IPTG gels exhibited high monotonicity scores, indicating a net decrease of modulus across the gel. Lag-1 analysis quantifies how smoothly the G’ changes, by a correlation test to determine how different adjacent samples are from each other. Despite the 1000 µM IPTG gel having the same monotonicity as the 250 µM IPTG gel, their autocorrelation values highlight the stochasticity or smoothness of each respective condition. The 250 µM condition showed strong positive autocorrelation, indicating that neighboring regions had similar mechanical properties and that G’ changes gradually across the gel. The 1000 µM condition showed a weak negative autocorrelation, suggesting increased local variability and deviations from a smooth gradient despite overall higher crosslinking and a weak negative slope. While limited, the above analyses provide a framework for quantifying bulk hydrogel properties and more complex patterning systems. Finally, the results for both conditions are summarized as a heatmap in **Figure 4f**. These results show that biocatalyzed hydrogel stiffening can be tuned in response to not only the level of induction of the sender cells, but also the proximity to the sender cells. The data also demonstrate that higher induction does not always translate to more efficient patterning in 3D matrices. Since our materials system is diffusion-driven, it is important to control the level of induction to ensure that there is no further loss of resolution due to oversaturation. Together, these experiments highlight the potential of recombinant quorum sensing to translate positional information from diffuse signals into macroscopic changes in material properties such as modulus.

### Increased Genetic Logic and Computational Methods Lead to More Complex Patterns of Gene Expression

Having demonstrated the ability of sender-receiver strains to create patterned regions of gene expression, we evaluated the potential of our system to create more complex patterns. To this end, we developed a computational model using experimental data to simulate diffusion of OC6 from sender cell colonies and the resulting pattern of GFP expression. The model takes in the location and level of induction of the sender cells and produces a heatmap of OC6 concentration as well as one of GFP expression as determined by the fluorescence response curve from **Figure 1b**. **Figure 5b** validates the model’s ability to predict the size and shape of the pattern resulting from the intersection of OC6 gradients as calculated computationally (**Fig. 5a**). Varying the volume of sender cells plated for each OC6 source provides another handle for tuning the size and intensity of this composite pattern. Excitingly, the EET activity pattern also shows the expected shape, indicating modeling from GFP can be applied to EET (**Fig. 5c**).

**Figure 5.**
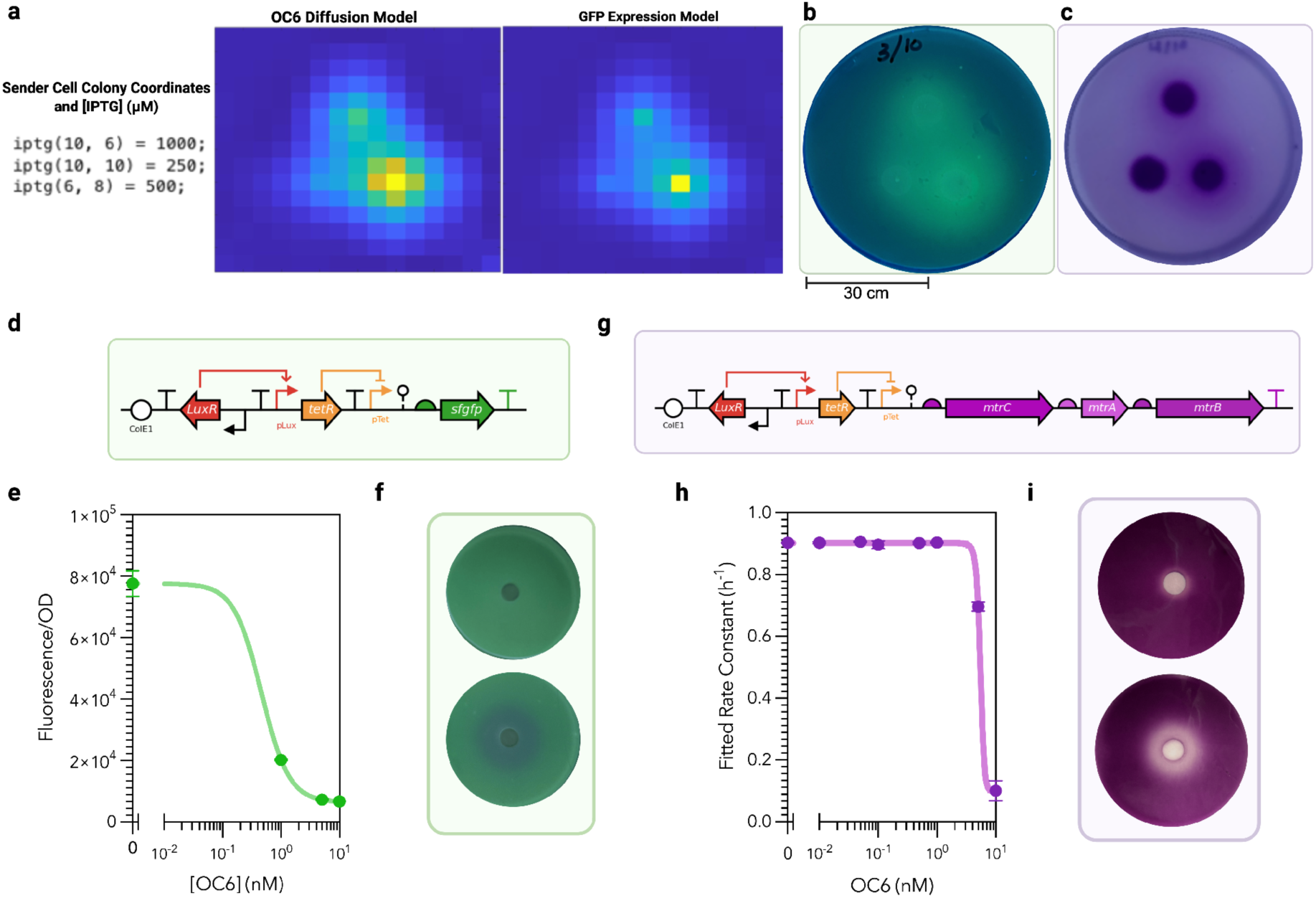
Increased genetic logic and computational methods leads to more complex patterns of gene expression. **a.**Computational model output of diffusion of OC6 from sender cell sources and the expected GFP response by the receiver cells. Included are the coordinates and levels of IPTG used as inputs for the model. **b.** Fluorescence plate assay recreating the sender cell placement from the computational model with 10 µL of sender cells at varying IPTG concentrations (250 µM, 500 µM, 1000 µM) per sender colony on a lawn of pAT3 receivers. **c.** Ferrozine plate assay recreating the sender cell placement from the computational model with 10 µL of sender cells per sender colony on a lawn of pAT4 receivers**. d.** Gene circuit diagram for the pAT3NOT plasmid controlling expression of sfgfp in *S.oneidensis* MR-1. **e.** pAT3NOT receiver cell OC6-HSL fluorescence response function. **f.** Patterned gene “turn-off” behavior shows regions of fluorescence only outside of the OC6 diffusion gradient which increases in size depending on the IPTG concentration in the sender cell colony (top: uninduced ; bottom: 1 mM IPTG). **g.** Gene circuit diagram for the pAT4NOT plasmid controlling expression of the mtrCAB operon in *S.oneidensis* JG1194. **h.** pAT4NOT receiver cell OC6-HSL fluorescence response function. **i.** Patterned gene “turn-off” behavior shows regions of Fe(III) reduction only outside of the OC6 diffusion gradient which increases in size depending on the IPTG concentration in the sender cell colony (top: uninduced ; bottom: 1 mM IPTG). Response curves represent the mean ± SEM of the appropriate measurement for *n* = 3 biological replicates. Scale bar = 30 mm.

Another method of developing more complex patterns is to increase the level of computation performed by the receiver cells. That is, have them interpret the induction level by regulating a cascade of activators and repressors much in the way biological systems react to positional chemical stimuli^18^. To this end, the pAT3 receiver cell plasmid was modified to exhibit NOT behavior resulting in a “turn-off” response rather than a “turn-on” response to OC6 as was seen in the original buffer gates. The OC6-activated LuxR now activates transcription of the TetR repressor protein downstream of the *pLux* promoter. The genetic output of the plasmid is now downstream of the P*_Tet_* promoter, resulting in repression of the constitutively expressed output protein only when OC6 is present to activate the synthesis of the repressor **(Fig. 5d, S12).**

First, we confirmed the expected negative Hill function behavior of the pAT3NOT circuit under variable induction conditions through fluorescence and ferrozine assays with lab stocks of OC6. The system showed the expected decrease in gene expression as OC6 concentration increased **(Fig. 5e).** Next, we set up the two-dimensional pattern plate assay with the NOT receiver cells which resulted in patterns inverse of its buffer gate counterpart **(Fig 5f).** Again, the size of the pattern can be modulated by changing the volume or level of induction of the sender cells **(Fig. S12).** The pAT4NOT plasmid uses the same regulatory architecture to control “turn-off” behavior of constitutively expressed MtrCAB. The pattern resulting from the two-dimensional plate assay was not as robust as the GFP “turn-off” behavior in response to what should be OC6 gradients that are similar in size. This can be explained by the long half-life of the MtrCAB complex in the *S. oneidensis* membrane. Before induction with OC6 the pAT4NOT plasmid is constitutively transcribing copies of MtrCAB which saturates the membrane and is stable even after transcription is halted by the activated TetR. For this reason, cells which have stopped producing MtrCAB may still exhibit electroactivity through existing copies of the complex. This issue could be addressed through lowering the level of constitutive expression through promoter or RBS engineering. Overall, these data demonstrate the wide variety of patterns possible with even the most basic of cell-cell communication circuits (**Fig. S14)**.

## Discussion and Conclusions

Here we establish a tunable system that links quorum sensing-mediated transcriptional regulation in *S.oneidensis* to macroscopic material properties. We advance synthetic biology tools in *Shewanella oneidensis* through the development of a modular recombinant quorum sensing system whose architecture is primed for adaptation of other quorum sensing signals beyond the LuxIR system^82^. We also expand the complexity of engineered living materials by leveraging engineered intercellular communication to control gene expression and chemical conversions using synthetic consortia. Sender-receiver communication enables tunable control over gene expression through both induction level and population composition. Synthetic consortia have been previously regulated using quorum sensing to maintain strain ratios or to share the metabolic burden of biosynthesis of industrially relevant molecules^83,84^. We expand on and apply a similar strategy to *S.oneidensis* which can be used to expand the substrate scope and synthetic capabilities of bacterial consortia.

Most importantly, this system recapitulates two-dimensional pattern formation through positional interpretation of autoinducer diffusion gradients. Direct control over material formation more faithfully captures the principles of morphogenesis, where diffuse chemical signals are interpreted to generate spatially organized physical structures. Embedding engineered “sender” and “receiver” *S. oneidensis* strains within hydrogel precursors and using autoinducer signals to control EET gene expression enables spatiotemporal interpretation of chemical signals to directly drive material formation. Whereas most synthetic patterning systems rely on fluorescence as the sole output, our approach couples spatial gene expression patterns to iron reduction, CuAAC chemistry, and hydrogel crosslinking through EET in *S.oneidensis*. These results support a design-test-build workflow where fluorescence responses of genetic circuits, patterning or otherwise, can be used to iterate on, visualize, and optimize cell responses and serves as proof of concept of the ease of adapting existing circuits to control biocatalysis of biorthogonal chemistry and material formation and properties^73^.

Material responses in the form of hydrogel crosslinking and CuAAC biocatalysis can be spatiotemporally regulated as a function of distance from the autoinducer source. The patterned hydrogel system establishes proof of principle for biologically programmed material differentiation. Importantly, the resulting differences in stiffness arose from the interactions between signal diffusion, receiver genetic logic, and localized. Lag-1 autocorrelation and monotonicity scores of patterned hydrogel strips provide insight into system behavior and stochasticity as a result of the dynamic induction from sender cell populations. Our analyses suggest that sender induction controls not only the magnitude of hydrogel stiffening, but also the extent to which spatial information is preserved and translated into patterned material properties. Further experiments are needed to examine the consistency of these crosslinking patterns and our ability to modulate the spatial pattern of crosslinking using sender cell induction and placement.

The patterned crosslinking of the hydrogel precursor is especially promising for biocompatible applications because hydrogel mechanical properties influence cell morphology and differentiation from a pluripotent state^85^. Further investigations include determining the smallest “resolution” possible with patterned crosslinking by using nanoindentation or AFM measurements to measure crosslinking around individual sender and receiver colonies. Importantly, our system is also amenable to more complex genetic logic than the standard activating buffer gate. NOT genetic in receiver results in inverse patterns of gene expression where OC6 acts as an inhibitory signal. Combining receiver cells with different genetic responses to sender cell signaling expands the complexity of patterns by multiplexing the interpretation of a single spatial signal. This lays the groundwork for further recapitulation of developmental systems by engineering regulatory mechanisms in the receiver cells to interpret multiple diffuse signals in activating and inhibitory capacities to create nested regions of gene expression. When applied to materials, this bodes well for subtractive rather than additive material formation and even patterned “differentiation” of more than one material state by tuning EET responses beyond on/off behaviors. Future work may incorporate more complex logic such as Boolean (AND, NAND, OR, NOR) computations and more than one inducer. By expanding the complexity of genetic logic, adding more experimental data to the model, and combining receiver strains, we are primed to use a build-test-learn approaches to program complex pattern formation applicable to orthogonal biocompatible chemistries including the formation of living hydrogels.

Overall, we present a framework for programming spatiotemporal differentiation of living material properties through quorum sensing-mediated control of EET activity. Our work enables the design of materials with spatially encoded functionality, with potential applications in biosensing in animal or environmental systems, localized drug delivery, and creating patterned scaffolds for tissue engineering. This system lays the groundwork at the intersection of synthetic biology, engineered living materials, redox chemistry, and developmental biology to create programmable materials that exhibit life-like properties including self-organization and patterned regions of differentiation.

## Materials and Methods

### Materials

4-arm 5k poly(ethylene glycol) methacrylate (PEG-MA, ≥95% functionalization, Advanced BioChemicals), copper(II) bromide (CuBr_2_, Sigma-Aldrich, 99%), tris(2-pyridylmethyl)amine (TPMA, Sigma-Aldrich, 98%), Tris(benzyltriazolylmethyl)amine (THPTA, Sigma-Aldrich, 95%), 2-(4-((bis((1-(tert-butyl)-1H-1,2,3-triazol-4-yl)methyl)amino)methyl)-1H-1,2,3-triazol-1-yl)acetic acid (BTTAA, Click Chemistry Tools > 95%), 2-hydroxyethyl 2-bromoisobutyrate (HEBIB, Sigma-Aldrich, 95%), sodium DL-lactate (NaC_3_H_5_O_3_, TCI, 60% in water), sodium fumarate (Na_2_C_4_H_2_O_4_, VWR, 98%), HEPES buffer solution (C_8_H_18_N2O_4_S, VWR, 1 M in water, pH = 7.3), potassium phosphate dibasic (K_2_HPO_4_, Sigma-Aldrich), potassium phosphate monobasic (KH_2_PO_4_, Sigma-Aldrich), sodium chloride (NaCl, VWR), ammonium sulfate ((NH_4_)_2_SO4, Fisher Scientific), magnesium(II) sulfate heptahydrate (MgSO_4_·7H2O, VWR), trace mineral supplement (ATCC), casamino acids (VWR), isopropyl ß-D-1-thiogalactopyranoside (IPTG, Teknova), 3-oxohexanoyl-homoserine lactone (OC6, Sigma-Aldrich), kanamycin sulfate (C_18_H_38_N_4_O_15_S, Growcells), iron(III) citrate (C_6_H_5_FeO_7,_ Alfa Aesar), iron (II) sulfate heptahydrate (FeSO_4_ · 7H_2_O, Alfa Aesar), 3-(2-Pyridyl)-5,6-bis(4-sulfophenyl)-1,2,4-triazine disodium salt hydrate (ferrozine, C_20_H_12_N_4_Na_2_O_6_S_2_, TCI), 5-norbornene-2-carboxylic acid (Sigma-Aldrich), agarose (C_10_H_15_N_3_O_3_, VWR), acryloyl chloride (C_3_H_3_ClO, Sigma-Aldrich), CalFluor 488 (Click Chemistry Tools), alkyne-PEG_4_-acid (Click Chemistry Tools) were used as received. Two-part silicone elastomer (SylgardTM 184, Electron Microscopy Sciences) was used according to manufacturer instructions. All media components were autoclaved or sterilized using 0.22 μm PES filters.

### Bacterial Strains and Culture

Bacterial strains and plasmids are listed in Table S3. Cultures were prepared as follows: bacterial stocks stored in 20% glycerol at −80 °C were streaked onto LB agar plates (for wild-type and knockout strains) or LB agar with 20 or 25 μg/mL kanamycin (for plasmid-harboring strains) and grown overnight at 30 °C for Shewanella and 37 °C for E. coli. Single colonies were isolated and inoculated into Shewanella Basal Medium (SBM) supplemented with 100 mM HEPES, 0.05% trace mineral supplement, 0.05% casamino acids, and 20 mM sodium lactate (2.85 μL of 60% w/w sodium lactate per 1 mL culture) as the electron donor. Aerobic cultures were pregrown in 15 mL culture tubes at 30 °C and 250 rpm shaking. Anaerobic cultures were pregrown using the same procedure outlined above, but in degassed growth medium in a humidified anaerobic chamber (3% H2, balance N2, Coy) and supplemented with 40 mM sodium fumarate (40 μL of 1 M sodium fumarate per 1 mL culture) as the electron acceptor. For stationary phase conditions, inducible strains were pregrown anaerobically without inducer(s) for 4–6 h before being diluted 1:25 into inducer-containing media (from 1000x stocks) to grow overnight. Cultures were washed 3x after pregrowth using SBM supplemented with 0.05% casamino acids (degassed for anaerobic cultures). Absorbance at 600 nm was measured using a BMG LABTECH CLARIOstar plate reader.

### Plasmid Construction

All bacterial strains, plasmids, genetic circuit maps, and sequence information for each genetic part are detailed in Tables C.1 and C.2. All plasmids were purchased from Addgene or assembled via Golden Gate cloning procedures using enzymes (BsaI, SapI, BsmBI) and buffers from New England Biolabs. DNA fragments used in Golden Gate cloning were generated via partial/whole-plasmid PCR or commercially synthesized (Integrated DNA Technologies or Twist Biosciences). Generally, 10 μL Golden Gate reactions were set up that contained 5 fmol of plasmid backbone and 20 fmol of each synthesized gene and/or PCR insert (as necessary). In a thermocycler, Golden Gate reactions were cycled 25-45 times, depending on the complexity and size of the construct: 90 s at 37 °C (for BsaI and SapI) or 42 °C (for BsmBI) followed by 3 min at 16 °C. After the cycles, reactions were incubated at 37 °C (for BsaI and SapI) or 55 °C (for BsmBI) for 5 min, 80 °C for 10 min, and then held at 4 °C. Golden Gate reactions were used to directly transform freshly prepared electrocompetent S. oneidensis or chemically competent E. coli strains. To prepare electrocompetent S. oneidensis, 5 mL of overnight S. oneidensis growth in LB medium at 30 °C was washed 3 times with sterile 10% glycerol at room temperature and concentrated to ∼300 μL. A 2 μL portion of Golden Gate reaction was mixed with 30 μL of concentrated electrocompetent S. oneidensis, transferred to a 1 mm electroporation cuvette, and electroporated at 1250 V. To recover electroporated cells, 250 μL of LB warmed in a 30 °C incubator was immediately added post-electroporation and cells were incubated/shaken at 30 °C and 250 rpm. After 2 h of recovery, 100 μL of cell suspension was plated onto LB agar plates containing 20 or 25 μg • mL-1 kanamycin and incubated overnight at 30 °C to obtain single colonies. Single colonies were used to inoculate LB liquid medium containing 20 or 25 μg • mL-1 kanamycin sulfate and 3 incubated/shaken overnight at 30 °C and 250 rpm. 5-alpha Competent E.coli (NEB) were used as directed for transformation into E.coli. These cultures were used to generate 20–22.5% glycerol stocks which were stored at −80 °C, and to harvest assembled plasmid for whole plasmid nanopore sequencing (Plasmidsaurus).

### Functional Verification of MtrC Expression

Strains containing mtrC expression circuits were functionally validated using an in situ Fe(III) reduction/ferrozine assay as previously described^1^. Briefly, strains were aerobically pregrown for ca. 3 h in SBM containing 20 mM lactate, 40 mM fumarate, and 20 or 25 µg • mL-1 kanamycin depending on the strain at 30°C with shaking at 250 rpm. These cell suspensions were diluted 25-fold into SBM containing 20 mM lactate, 40 mM fumarate, kanamycin, and appropriate inducers, sealed with impermeable foil (Thermo Scientific) then allowed to grow for ca. 18 h at 30°C. Subsequently, these growths were diluted 250-fold into 96-well plates containing SBM solution with 20 mM lactate, 20 or 25 µg • mL-1 kanamycin, 1 mg • mL-1 ferrozine, appropriate inducers, and 5 mM Fe(III) citrate, such that the final well volume was 250 µL. Fe(II) standards were also included in the plate using dissolved FeSO_4_. The 96-well plate was sealed with a sterile/optically transparent film (PCR-SP-S, AxySeal Scientific), covered with a polystyrene plate lid (Eppendorf) with silicone grease lining the edges, removed from the anaerobic chamber, and placed within a BMG LABTECH CLARIOstar plate reader with temperature control set to 30 °C. Without shaking, the absorbance at 562 nm was measured every 10 min for at least 14 h. Using the Fe(II) standards, raw kinetics data was converted to Fe(II) concentrations vs. time. Fe(II) kinetics for individual replicates were background subtracted (i.e. Fe(II) level at the initial time point) and fitted to an exponential Monod-type model to obtain fitted rate constants (μ):
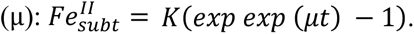.

### Quantification and Modeling of Fluorescence Constructs

Strains containing sfgfp architectures were aerobically pregrown overnight using standard culture conditions outlined above, with the addition of 20 or 25 µg • mL-1 kanamycin depending on the strain. Cultures were then diluted 25-fold into 96-well plates containing SBM with 0.05% casamino acids, 20 mM lactate, 40 mM fumarate, kanamycin, and varying amounts of inducer(s) (from 25X stocks). Plates were covered with a polystyrene plate lid (Eppendorf) and placed at 30°C for 18–24 h. sfGFP fluorescence (488/530 nm) and cell suspension absorbance (600 nm) were measured using a BMG LABTECH CLARIOstar plate reader to yield fluorescence • absorbance−1 for each sample. For each sample, the background fluorescence • absorbance−1 from an empty vector (pCD8) control was subtracted. In addition, strains were normalized in each plate to a RNAP flux standard strain constitutively expressing sfgfp (pCDe1) via the Ptrc* promoter to enable relative expression unit (REU) calculations. A nonlinear fitting algorithm in GraphPad Prism 9 was used to fit inducible gene expression to the following activating Hill function: *y* = min + (max − min) [*I*] *n K*1/2 *n*+ [*I*]*n*. Further details on modeling can be found in previous work^1,2^. Fitting parameters and “goodness of fit” can be found in Table S3.

### Fluorescence Protein Pattern Assay

Sender and fluorescent receiver strains in S.oneidensis were aerobically pregrown overnight using standard culture conditions outlined above, with the addition of 25 µg • mL-1 kanamycin. Cultures were washed 3x after pregrowth using SBM supplemented with 0.05% casamino acids (SBM+cas). Absorbance at 600 nm was measured using a BMG LABTECH CLARIOstar plate reader. Receiver cells were diluted to an OD600 of 0.15 in 0.7% agarose in SBM+cas and spread evenly on a 60 mm 1.5% SBM + cas agarose plate supplemented with 20 mM sodium lactate, 40 mM sodium fumarate, and 25 µg • mL-1 kanamycin. Sender cells were diluted to an OD600 of 0.3 in 1 mL and then concentrated 50-fold in SBM+cas supplemented with 20 mM sodium lactate, 40 mM sodium fumarate, 25 µg • mL-1 kanamycin, and variable amounts of IPTG as inducer. 20 µL of the concentrated sender cells were pipetted onto a 6 mm Whatman filter paper disk and placed in the center of the receiver top agar. The plates were incubated aerobically for 18 h at 30° C. The fluorescent protein expression pattern was visualized and imaged using an UV Transilluminator (insert brand).

### EET Protein Pattern Assay

Sender and EET-protein-expressing receiver strains in S.oneidensis were aerobically pregrown overnight using standard culture conditions outlined above, with the addition of 25 µg • mL-1 kanamycin. Cultures were washed 3x after pregrowth using SBM supplemented with 0.05% casamino acids (SBM+cas). Absorbance at 600 nm was measured using a BMG LABTECH CLARIOstar plate reader. Receiver cells were diluted to an OD600 of 0.15 in 0.7% agarose in SBM+cas and spread evenly on a 60 mm 1.5% SBM + cas agarose plate supplemented with 20 mM sodium lactate, 40 mM sodium fumarate, and 25 µg • mL-1 kanamycin. Sender cells were diluted to an OD600 of 0.3 in 1 mL and then concentrated 50-fold in SBM+cas supplemented with 20 mM sodium lactate, 40 mM sodium fumarate, 25 µg • mL-1 kanamycin, and variable amounts of IPTG as inducer. 20 µL of the concentrated sender cells were pipetted onto a 6 mm Whatman filter paper disk and placed in the center of the receiver top agar. The plates were incubated aerobically for 18 h at 30° C. The pattern of EET activity was measured by pouring over 5 mL of 0.7% agarose in SBM+cas containing 2 mM Fe(III) and 1 mM ferrozine over each plate followed by a 10 minute incubation. The EET activity pattern was visualized and imaged using a white light box (insert brand).

### CuAAC Reaction Pattern Assay

Sender and EET-protein-expressing receiver strains in S.oneidensis were aerobically pregrown overnight using standard culture conditions outlined above, with the addition of 25 µg • mL-1 kanamycin. Cultures were washed 3x after pregrowth using SBM supplemented with 0.05% casamino acids (SBM+cas). Absorbance at 600 nm was measured using a BMG LABTECH CLARIOstar plate reader. Receiver cells were diluted to an OD600 of 0.15 in 0.7% agarose in SBM+cas and spread evenly on a 60 mm 1.5% SBM + cas agarose plate supplemented with 20 mM sodium lactate, 40 mM sodium fumarate, and 25 µg • mL-1 kanamycin. Sender cells were diluted to an OD600 of 0.3 in 1 mL and then concentrated 50-fold in SBM+cas supplemented with 20 mM sodium lactate, 40 mM sodium fumarate, 25 µg • mL-1 kanamycin, and variable amounts of IPTG as inducer. 20 µL of the concentrated sender cells were pipetted onto a 6 mm Whatman filter paper disk and placed in the center of the receiver top agar. The pattern of EET activity was measured by pouring 3 mL of 0.7% agarose in SBM+cas containing 100 µM alkyne-PEG_4_-acid, .6 µM CalFluor488, 400 µM Cu:BTTAA (1:6), and 12 mM fumarate over each plate followed by a 1 h anaerobic incubation. The successful formation of the CuAAC fluorescent probe was visualized and imaged using a UV transilluminator (insert brand).

### Hydrogel Pattern Assembly

A PDMS mold was prepared by pouring the liquid polymer into a rectangular mold made from microscope slides. The bottom of the mold contained a .5 mm tall, 65 mm long, and 10 mm wide silicone spacer. Following a 4 hour cure at 60°C, the PDMS mold was removed from the glass enclosure and the silicone spacer was removed from the now solid yet flexible hydrogel mold leaving a narrow channel for casting the hydrogel precursor containing receiver cells. The final concentrations in the precursor solution were 1 wt % ACAG, 5 wt % PEG-MA, 10 μM Cu-TPMA, 100 μM HEBIB, 20 mM lactate, 40 mM fumarate, and 25 µg/mL kanamycin and receiver cells at an OD_600_ of 0.1. 500 µL of this mixture were dispensed into the PDMS mold before using a sterile cell spreader passed along the top of the channel to level out the hydrogel and ensure even thickness along the length of the gel. The 1% ACAG in the precursor was allowed to set at 4°C for 10 minutes before sealing the hydrogel, still in the mold, with a microscope slide on top. This assembly was sealed with a binder clip to prevent evaporation and the entire assembly was placed in an anaerobic chamber for overnight incubation at 30°C.

### Ferrozine Co-Culture Plate Assay

Sender and EET receiver strains were co-cultured in a ferrozine mixture as described above. Briefly, strains were individually aerobically pregrown overnight in SBM containing 20 mM lactate, 40 mM fumarate, and 20 or 25 µg • mL-1 kanamycin depending on the strains at 30°C with shaking at 250 rpm. Cultures were washed 3x after pregrowth using SBM supplemented with 0.05% casamino acids (SBM+cas). All cultures were normalized to an OD600 of 0.4. Then, sender and receiver cultures were mixed at varying sender:receiver ratios. Subsequently, these co-cuture mixtures were diluted anaerobically 25-fold into 96-well plates containing SBM solution with 20 mM lactate, 20 or 25 µg • mL-1 kanamycin, 1 mg • mL-1 ferrozine, 1 mM IPTG as an inducer for the sender cells, and 5 mM Fe(III) citrate, such that the final well volume was 250 µL. Fe(II) standards were also included in the plate using dissolved FeSO_4_. The 96-well plate was sealed with a sterile/optically transparent film (PCR-SP-S, AxySeal Scientific), covered with a polystyrene plate lid (Eppendorf) with silicone grease lining the edges, removed from the anaerobic chamber, and placed within a BMG LABTECH CLARIOstar plate reader with temperature control set to 30 °C. Without shaking, the absorbance at 562 nm was measured every 10 min for at least 14 h. Using the Fe(II) standards, raw kinetics data was converted to Fe(II) concentrations vs. time. Fe(II) kinetics for individual replicates were background subtracted (i.e. Fe(II) level at the initial time point) and fitted to an exponential Monod-type model to obtain fitted rate constants (μ): 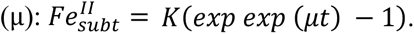.

The fitted rate constants were then plotted as a heatmap in GraphPad Prism 10.

### Fluorescence Co-Culture Plate Assay

Sender and fluorescent receiver strains were aerobically pregrown overnight using standard culture conditions outlined above, with the addition of 20 or 25 µg • mL–1 kanamycin depending on the strain. Cultures were washed 3x after pregrowth using SBM supplemented with 0.05% casamino acids (SBM+cas). All cultures were normalized to an OD600 of 0.4. Cultures were then diluted 25-fold into 96-well plates containing SBM with 0.05% casamino acids, 20 mM lactate, 40 mM fumarate, kanamycin, and varying amounts of IPTG (from 25X stocks). Plates were covered with a polystyrene plate lid (Eppendorf) and placed at 30 °C for 18–24 h. sfGFP fluorescence (488/530 nm) and cell suspension absorbance (600 nm) were measured using a BMG LABTECH CLARIOstar plate reader to yield fluorescence • absorbance−1 for each sample. For each sample, the background fluorescence • absorbance−1 from an empty vector (pCD8) control was subtracted. A nonlinear fitting algorithm in GraphPad Prism 9 was used to fit inducible gene expression to the following activating Hill function: *y* = min + (max − min) [*I*] *n K*1/2 *n*+ [*I*]*n*. Further details on modeling can be found in previous work^1,2^. Fitting parameters and “goodness of fit” can be found in Table S3. The endpoint fluorescence values normalized by cell density were plotted as a heatmap in GraphPad Prism 10.

### CuAAC Co-Culture Plate Assay

All reactions were performed in *Shewanella* basal media (SBM) supplemented with 0.05% w/v casamino acids with lactate (20 mM) as a carbon source and fumarate (20 mM) as the primary electron acceptor. Stock solutions of 1 M sodium fumarate and 60 w/v% lactate solutions were stored at 4 °C until use. Aliquots of 1.2 mM of CalFluor 488 were created in DMSO and stored frozen at −80 °C until use. Aliquots of 4 mM alkyne-PEG_4_-acid were created in DMSO and stored at −20 °C until use. An 8 mM copper bromide stock in sterile water was created and stored at 4 °C and mixed with an equal volume amount of 48 mM freshly made stock of BTTAA in sterile water. Sender and EET receiver strains were individually aerobically pregrown overnight in SBM containing 20 mM lactate, 40 mM fumarate, and 20 or 25 µg • mL–1 kanamycin depending on the strains at 30°C with shaking at 250 rpm. Cultures were washed 3x after pregrowth using SBM supplemented with 0.05% casamino acids (SBM+cas). All cultures were normalized to an OD600 of 0.4. Then, sender and receiver cultures were mixed at varying sender:receiver ratios.In order, the following was added to yield a 200 μL reaction in either degassed or ambient SBM supplemented with 0.05% casamino acids with the final concentrations: lactate (20 mM), fumarate (20 mM), Cu:BTTAA 1:6 (8 μM:48 μM), alkyne-PEG_4_-acid (100 μM), CalFluor 488 (0.6 μM), and finally *S. oneidensis* (OD_600_ of 0.2). Fluorescence emission was collected on a BMG LABTECH CLARIOstar plate reader with a 491 (±14) nm and an emission collection at 538 (±38) nm). After the addition of all plate components, the 96-well plate was sealed with a sterile and optically transparent sealing film (PCR-SP-S, AxySeal Scientific) and covered with a polystyrene plate lid (Eppendorf) lined with silicone grease and sealed with Teflon tape. The plate reader was held at 30 °C and collected emissions every 90 s for 14h. Fluorescence measurements were normalized by OD600 and fit using nonlinear regression fit to a second order polynomial equation Y=B0 + B1*X + B2*X^2 in GraphPad Prism 10. The B2 values were plotted as a heatmap.

### Synthesis of Acrylated Agarose

1.0 g of agarose was dissolved in 25 mL of DMAc at 100 °C in a silicone oil bath. The agarose solution was then cooled to 0 °C in an ice-water bath, and 125µL of acryloyl chloride diluted in 1 mL DMAc was added slowly under stirring. The mixture was continuously stirred at 0 °C for 1 h and then at room temperature for 4 h. The product was precipitated and washed thoroughly by acetone and then dried in vacuum.

### Hydrogel Radical Cross-Linking Using Engineered *S.oneidensis*

CuBr_2_ and TPMA were dissolved at 8 mM in DMF for storage and combined into a 400 µM Cu-TPMA stock solution in DMF before reaction preparation. HEBIB (1.45 µL) was added to SBM with casamino acids (143 µL) to create a 69 mM stock solution which was diluted 5-fold in SBM with casamino acids to create a 13.8 mM solution before reaction preparation. Per 50 μL hydrogel disc that was analyzed by rheology, a cross-linking reaction mixture was prepared as follows: PEG-MA was dissolved at 6.18 wt % in SBM with 0.05% casamino acids and aliquoted into an autoclaved microfuge tube (40.47 μL). Solutions of 400 μM Cu-TPMA (0.625 μL or 1.25 µL), 13.8 mM HEBIB (0.3625 μL), 60% sodium lactate (0.143 μL), and 1 M sodium fumarate (2 μL) were added to the PEG-MA solution and mixed. For co-polymerization, ACAG was dissolved at 1.25 weight % in SBM with 0.05% casamino acids by placing on a heating block at 90°C with shaking. PEG-MA was then dissolved at 6.18 wt %. Per 50 μL gel mixture, the remaining 1.4 μL was used for antibiotic and inducing molecule addition where necessary. Constituent volumes were multiplied as necessary to create a single primary stock for each experiment involving identical inducer conditions and *S. oneidensis* strains. The final concentrations in solution were 1 wt % ACAG, 5 wt % PEG-MA, 10 μM Cu-TPMA, 100 μM HEBIB, 20 mM lactate, 40 mM fumarate, and 0, 20, or 25 µg/mL kanamycin where necessary, depending on *S. oneidensis* strain. Inducer concentrations ranged from 0 to 100 nM were diluted from 100x stock solutions. IPTG was dissolved in sterile H_2_O and OC6 was dissolved in DMF; all inducer stocks were stored at −20 °C. The primary gel mixture was then distributed into individual autoclaved microfuge tubes of 45 μL aliquots to which 5 μL of OD_600_-normalized cells were added. The gel solutions were mixed and dispensed onto hydrophobically treated glass slides with a 0.5 mm silicone spacer separating the two glass layers. The gels were allowed to react at 30 °C for 2 h at inoculating OD_600_ = 0.2. Hydrogels were removed from the slides using a razor blade and placed into 3 mL baths of 1x PBS overnight to swell to equilibrium at room temperature in the dark.

### Rheological Analysis

Swollen hydrogels prepared as outlined above were analyzed by oscillatory shear rheology using a TA Instruments Discovery HR-2 Rheometer utilizing Trios software with an 8 mm parallel plate geometry as outlined previously^55^. Briefly, the geometry gap was lowered to an axial force at or above 0.02 N (usually between 300−600 μm, depending on the cross-link density and swelling ratio). Storage was measured using frequency sweeps from 0.1 to 1 Hz at a constant strain of 1%. Moduli for a single gel were quantified by averaging the linear viscoelastic region of each frequency sweep.

### Computational Model

Three models were iteratively developed in MATLAB’s app designer. The first two models, mps_v1.mlapp and mps_v2.mlapp, use an approximation for the AHL surface diffusion constant based on the time required to diffuse to the edge of a 145 cm^2^ culture plate using penetration theory. To model expression of LuxI from IPTG, OC6 from LuxI, and GFP from OC6, standard Hill functions were used. First order chemical decay was assumed. Expression and decay were combined to produce chemical equations of the form below, where X is the concentration of LuxI, OC6, or GFP. Extracellular diffusion was only modeled for OC6. Each model utilizes MATLAB’s lsqnonlin function, part of the Optimization Toolbox add-in, to numerically solve the chemical equations for steady state concentrations. In the code, the equations are contained in the *patterning* function with lsqnonlin implemented in the *model* function.

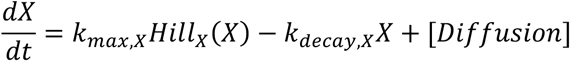

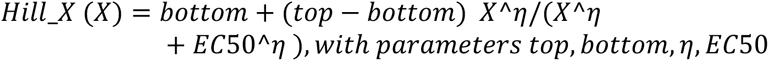

mps_v1.mlapp estimates all Hill parameters and decay constants from educated guesses. mps_v2.mlapp models OC6 expression directly from IPTG using experimental Hill parameters. mps_v2.mlapp models GFP steady state REU from OC6 using experimental Hill parameters from a pAT3 curve. Decay constants and weighting of expression vs decay based on *k_max_* were tuned to match experimental measurements of 0.9 REU GFP expression after 18 hours. As a result, mps_v2.mlapp only used one chemical equation for OC6 expression, diffusion, and decay.

The final model, mps.m, utilized many of the same expression and decay parameters as mps_v2.mlapp. Two parameters were modified based on experiment, however. The OC6 decay constant was referenced from Horswill et al.,^86^ and the OC6 diffusion constant on agar was computed using the diffusion constant for OC12 in water from Stewart^87^ reduced by a factor of 0.9 based on Fatin-Rouge et al.’s analysis of agar effects on diffusion^88^.

Currently, the concentration of OC6 at the sender cells is set to the steady state value in mps_v2.mlapp and mps.m. This assumes that OC6 is always diffusing away from sender cells, which is not always the case. This could be improved by modeling the actual steady state expression, decay, and diffusion from sender cells which could better capture cases where a highly expressing sender cell causes OC6 to diffuse towards another adjacent sender cell.

These models were converted into MATLAB apps with user interface and usage detailed in the accompanying presentation Shew_Patterning_Model-final.pptx. Within the *compute_GFP* function, all apps currently include two-input genetic circuit logic (AND, OR, NAND, NOR) for receiver cell GFP expression. However, the actual models for how receiver cells respond to varying input concentrations have not been developed, and require future computational literature review and experiments.

## Supporting Information

The following files are available free of charge. SupplementaryInformation.pdf

## Author Contributions

The manuscript was written through contributions of all authors. All authors have given approval to the final version of the manuscript. I.E.M., G.P., and B.K.K. conceived the project and designed research. I.E.M., V.S.D., and G.P. performed crosslinking experiments and rheological analysis.

A.Y.L. developed the computational model. I.E.M. performed cloning and circuit characterization by growth, fluorescence, iron reduction assays, and biocatalyzed CuAAC. I.E.M. performed co-culture assays, plate patterning assays, image analysis and model validation.

## Acknowledgements

We would like to thank Prof. Jeffrey Gralnick (U. Minnesota) for generously providing *S. oneidensis* strains Δ*mtrC*/Δ*omcA*/Δ*mtrF* and ΔmtrC*/*ΔomcA/ΔmtrF/ΔmtrA/ΔmtrD/ΔdmsE/ ΔSO4360/ΔcctA/ ΔrecA. This research was financially supported by the Welch Foundation (grant F-1929, B.K.K.), the National Institutes of Health under award number 1R35GM156283 (B.K.K.), and the Air Force Office of Scientific Research under award number FA9550-24-1-0273 (B.K.K.). We also acknowledge the use of shared research facilities supported in part by the Texas Materials Institute, the Center for Dynamics and Control of Materials: an NSF MRSEC (grant DMR-1720595). Schematics were created using BioRender. com, graphs were created in GraphPad Prism.

## Data Availability

Model code available at github.com/keitzlab. Raw data supporting the findings described in this manuscript can be accessed via the Texas Data Repository at https://doi.org/10.18738/T8/BQJ77V

