## Supplemental Information for "Quorum-Sensing-Mediated Extracellular Electron Transfer Enables Hydrogel Morphogenesis"

### Supplementary Information

**Table S1:** Bacterial strains and plasmids used in this study.

| ***S.oneidensis* Strains (+plasmid)** | **Description/Genotype** | **Reference or Source** | **Benchling Link** |
|---|---|---|---|
| MR-1 | MR-1 (ATCC700550), wild-type strain accession code NC_004347 | American-Type Culture Collection | |
| JG596 | *ΔmtrCΔomcAΔmtrF* | Jeffrey Gralnick, U. of Minnesota | |
| JG1194 (ΔMtr) | *ΔmtrC/ ΔomcA/ ΔmtrF/ ΔmtrA/ ΔmtrD/ ΔdmsE/ ΔSO4360/ ΔcctA/ ΔrecA* | Jeffrey Gralnick, U. of Minnesota | |
| MR-1 + pCDe1 | *sfgfp* REU plasmid ($P_{trc*}$) | [75] | pCDe1 |
| MR-1 + pCD8 | Empty buffer gate ($P_{tacsym0}$; LacI) | [75] | pCD8 |
| JG596 + pAT1 | *mtrC* buffer gate ($P_{lux}$; LuxR) | [75] | pAT1 |
| MR-1 + pAT3 | *sfgfp* buffer gate ($P_{lux}$; LuxR) | [75] | pAT3 |
| JG1196 +pAT4 | *mtrCAB* buffer gate ($P_{lux}$; LuxR) | This work | pAT4 |
| MR-1 + pCDh_LuxI | *luxI* buffer gate ($P_{tacsym0}$; LacI) | This work | pCDh_LuxI |
| JG596 + pCDh_LuxI | *luxI* buffer gate ($P_{tacsym0}$; LacI) | This work | |
| JG1194 + pCDh_LuxI | *luxI* buffer gate ($P_{tacsym0}$; LacI) | This work | |
| MR-1 + pAT3NOT | *sfgfp* NOT gate ($P_{lux}$; LuxR/$P_{tet}$; TetR) | This work | pAT3NOT |
| JG1194 + pAT4NOT | *MtrCAB* NOT gate ($P_{lux}$; LuxR/$P_{tet}$; TetR | This work | pAT4NOT |
| MR-1 + pAT3mScarlet | *mScarlet* buffer gate ($P_{lux}$; LuxR) | This Work | pAT3mScarlet |

**Table S2:** Genetic parts/sequences used to construct the plasmids in this study.

| Genetic Part | DNA Sequence (5' to 3') |
|---|---|
| **Promoters** | |
| $P_{tacsymO}$[88] | TGTTGACAATTAATCATCGGCTCGTATAATGTGTGGAATTGTGAGCGCT CACAATTCTATGGACTATGTTT |
| $P_{lacI}$ | GCGGCGCGCCATCGAATGGCGCAAAACCTTTCGCGGTATGGCATGATA GCGCCCGGAAGAGAG TCAATTCAGGGTGGTGAAT |
| $P_{tet}$[89] | TACTCCACCGTTGGCTTTTTTCCCTATCAGTGATAGAGATTGACATCCCT ATCAGTGATAGAGATA ATGAGCAC |
| $P_{luxB}$ | GCTTAACGATCGTTGGCTGACCTGTAGGATCGTACAGGTTTACGCAAGA AAATGGTTTGTTACAG TCGAATAAA |
| $P_{trc*}$[90] | GAAATATTCTGAAATGAGCTGTTGACAATTAATCATCCGGCTCGTATAAT GTGTGGAA |
| **Ribosome Binding Sites** | |
| B0032 | TCACACAGGAAAGTACTAG |
| RBS lux1 | GGAAGAGAGTCAATTCAGGGTGGTGAAT |
| tetR RBS | CTATGGACTATGTTTTCACACAGGAAAGGCCTCG |
| sRBS1mtrC[75] | GGGGAAAAACAGCAGTGCGAT |
| sRBS1_pCDf1mtrA[75] | ATAGGCGGCTTCATTGACGGTCCCA |
| **Ribozymes** | |
| RiboJ[91] | AGCTGTCACCGGATGTGCTTTCCGGTCTGATGAGTCCGTGAGGACGAA ACAGCCTCTACAAATAATTTTGTTTAA |
| **Terminators** | |
| B0015 | TAATCTAGACCAGGCATCAAATAAAACGAAAGGCTCAGTCGAAAGACT GGGCCTTTCGTTTTATCTGTTGTTTGTCGGTGAACGCTCTCTACTAGAG TCACACTGGCTCACCTTCGGGTGGGCCTTTCTGC GTTTATA |

| | |
|---|---|
| T0 | CTTGGACTCCTGTTGATAGATCCAGTAATGACCTCAGAACTCCATCTGG<br>ATTTGTTCAGAACGCTC<br>GGTTGCCGCCGGGCGTTTTTTATTGGTGAGAATCCAAGCA |
| ECK120029600[92] | TTCAGCCAAAAAACTTAAGACCGCCGGTCTTGTCCACTACCTTGCAGT<br>AATGCGGTGGACAGGAT CGGCGGTTTTCTTTTCTCTTCTCAA |
| ECK120017009[92] | GATCTAACTAAAAAGGCCGCTCTGCGGCCTTTTTTCTTTTCACT |
| ECK120033736[92] | AACGCATGAGAAAGCCCCCGGAAGATCACCTTCCGGGGGCTTTTTTAT<br>TGCGC |
| **Genes** | |
| *luxI* | ATGACTATAATGATAAAAAAATCGGATTTTTTGGCAATTCCATCGGAGG<br>AGTATAAAGGTATTCTAAGTCTTCGTTATCAAGTGTTTAAGCAAAGACT<br>TGAGTGGGACTTAGTTGTAGAAAATAACCTTGAATCAGATGAGTATGAT<br>AACTCAAATGCAGAATATATTTATGCTTGTGATGATACTGAAAATGTAAG<br>TGGATGCTGGCGTTTATTACCTACAACAGGTGATTATATGCTGAAAAGT<br>GTTTTTCCTGAATTGCTTGGTCAACAGAGTGCTCCCAAAGATCCTAATA<br>TAGTCGAATTAAGTCGTTTTGCTGTAGGTAAAAATAGCTCAAAGATAAA<br>TAACTCTGCTAGTGAAATTACAATGAAACTATTTGAAGCTATATATAAAC<br>ACGCTGTTAGTCAAGGTATTACAGAATATGTAACAGTAACATCAACAGC<br>AATAGAGCGATTTTTAAAGCGTATTAAAGTTCCTTGTCATCGTATTGGAG<br>ACAAAGAAATTCATGTATTAGGTGATACTAAATCGGTTGTATTGTCTATG<br>CCTATTAATGAACAGTTTAAAAAAGCAGTCTTAAATTAA |
| *lacI* | GTGAAACCAGTAACGTTATACGATGTCGCAGAGTATGCCGGTGTCTCTT<br>ATCAGACCGTTTCCCG<br>CGTGGTGAACCAGGCCAGCCACGTTTCTGCGAAAACGCGGGAAAAAG<br>TGGAAGCGGCGATGGCGGAGCTGAATTACATTCCCAACCGCGTGGCAC<br>AACAACTGGCGGGCAAACAGTCGTTGCTGATT<br>GGCGTTGCCACCTCCAGTCTGGCCCTGCACGCGCCGTCGCAAATTGTC<br>GCGGCGATTAAATCTC<br>GCGCCGATCAACTGGGTGCCAGCGTGGTGGTGTCGATGGTAGAACGAA<br>GCGGCGTCGAAGCCT<br>GTAAAGCGGCGGTGCACAATCTTCTCGCGCAACGCGTCAGTGGGCTGA<br>TCATTAACTATCCGCT<br>GGATGACCAGGATGCCATTGCTGTGGAAGCTGCCTGCACTAATGTTCC<br>GGCGTTATTTCTTGATG<br>TCTCTGACCAGACACCCATCAACAGTATTATTTTCTCCCATGAAGACGG<br>TACGCGACTGGGCGTG |

| | |
|---|---|
| | GAGCATCTGGTCGCATTGGGTCACCAGCAAATCGCGCTGTTAGCGGGC<br>CCATTAAGTTCTGTCT<br>CGGCGCGTCTGCGTCTGGCTGGCTGGCATAAATATCTCACTCGCAATCA<br>AATTCAGCCGATAGC<br>GGAACGGGAAGGCGACTGGAGTGCCATGTCCGGTTTTCAACAAACCAT<br>GCAAATGCTGAATGAG<br>GGTATCGTTCCCACTGCGATGCTGGTTGCCAACGATCAGATGGCGCTGG<br>GCGCAATGCGCGCC<br>ATTACCGAGTCCGGGCTGCGCGTTGGTGCGGATATCTCGGTAGTGGGAT<br>ACGACGATACCGAAG<br>ACAGCTCATGTTATATCCCGCCGTTAACCACCATCAAACAGGATTTTCG<br>CCTGCTGGGGCAAACC<br>AGCGTGGACCGCTTGCTGCAACTCTCTCAGGGCCAGGCGGTGAAGGG<br>CAATCAGCTGTTGCCC<br>GTGTCACTGGTGAAAAGAAAAACCACCCTGGCGCCCAATACGCAAAC<br>CGCCTCTCCCCGCGCGT<br>TGGCCGATTCATTAATGCAGCTGGCACGACAGGTTTCCCGACTGGAAA<br>GCGGGCAGTGA |
| *luxR* | ATGAAAAACATAAATGCCGACGACACATACAGAATAATTAATAAAATTA<br>AAGCTTGTAGAAGCAAT<br>AATGATATTAATCAATGCTTATCTGATATGACTAAAATGGTACATTGTGA<br>ATATTATTTACTCGCGA<br>TCATTTATCCTCATTCTATGGTTAAATCTGATATTTCAATCCTAGATAATTA<br>CCCTAAAAAATGGAG<br>GCAATATTATGATGACGCTAATTTAATAAAATATGATCCTATAGTAGATTA<br>TTCTAACTCCAATCATT<br>CACCAATTAATTGGAATATATTTGAAAACAATGCTGTAAATAAAAAATCT<br>CCAAATGTAATTAAAGA<br>AGCGAAAACATCAGGTCTTATCACTGGGTTTAGTTTCCCTATTCATACG<br>GCTAACAATGGCTTCGG<br>AATGCTTAGTTTTGCACATTCAGAAAAAGACAACTATATAGATAGTTTAT<br>TTTTACATGCGTGTATG<br>AACATACCATTAATTGTTCCTTCTCTAGTTGATAATTATCGAAAAATAAA<br>TATAGCAAATAATAAATC<br>AAACAACGATTTAACCAAAAGAGAAAAAGAATGTTTAGCGTGGGCATG<br>CGAAGGAAAAAGCTCTTGGGATATTTCAAAAATATTAGGTTGCAGTGA<br>GCGTACTGTCACTTTCCATTTAACCAATGCGCAAATGAAACTCAATACA<br>ACAAACCGCTGCCAAAGTATTTCTAAAGCAATTTTAACAGGAGCAATT<br>GATT GCCCATACTTTAAAAATTGA |

| | |
|---|---|
| *tetR* | ATGTCTCGTTTAGATAAATCTAAAGTTATCAACTCTGCTTTAGAATTATT<br>AAACGAAGTTGGTATCG<br>AAGGTTTAACTACTCGTAAATTAGCTCAAAAATTAGGTGTTGAACAGCC<br>CACATTATACTGGCACG<br>TTAAAAACAAGAGGGCGTTATTAGATGCTCTCGCTATCGAAATGTTAGA<br>TCGTCACCACACTCACT<br>TCTGTCCATTAGAAGGTGAATCTTGGCAAGATTTCTTACGTAACAACGC<br>TAAATCGTTCCGTTGTG<br>CGTTATTATCGCACCGTGATGGTGCTAAAGTTCACTTAGGTACTCGTCC<br>AACTGAAAAACAATACG<br>AAACTTTAGAAAACCAATTAGCTTTCTTATGTCAACAAGGTTTCTCGCT<br>CGAAAACGCGCTCTATG<br>CGTTATCGGCTGTTGGCCACTTCACTTTAGGTTGTGTTTTAGAAGATCA<br>AGAACACCAAGTTGCTA<br>AAGAAGAACGTGAAACTCCAACTACTGATTCTATGCCACCATTATTACG<br>TCAAGCTATCGAATTAT<br>TCGATCACCAAGGTGCCGAGCCAGCGTTCCTCTTCGGTTTAGAATTAAT<br>CATCTGTGGTTTAGAA AAACAATTAAAATGTGAATCTGGTTCTTAA |
| *sfgfp* | ATGCGTAAAGGCGAAGAGCTGTTCACTGGTGTCGTCCCTATTCTGGTG<br>GAACTGGATGGTGATG<br>TCAACGGTCATAAGTTTTCCGTGCGTGGCGAGGGTGAAGGTGACGCAA<br>CTAATGGTAAACTGAC<br>GCTGAAGTTCATCTGTACTACTGGTAAACTGCCGGTACCTTGGCCGACT<br>CTGGTAACGACGCTGA<br>CTTATGGTGTTCAGTGCTTTGCTCGTTATCCGGACCATATGAAGCAGCAT<br>GACTTCTTCAAGTCC<br>GCCATGCCGGAAGGCTATGTGCAGGAACGCACGATTTCCTTTAAGGAT<br>GACGGCACGTACAAAA<br>CGCGTGCGGAAGTGAAATTTGAAGGCGATACCCTGGTAAACCGCATTG<br>AGCTGAAAGGCATTGA<br>CTTTAAAGAAGACGGCAATATCCTGGGCCATAAGCTGGAATACAATTTT<br>AACAGCCACAATGTTTA<br>CATCACCGCCGATAAACAAAAAAATGGCATTAAAGCGAATTTTAAAATT<br>CGCCACAACGTGGAGG<br>ATGGCAGCGTGCAGCTGGCTGATCACTACCAGCAAAACACTCCAATCG<br>GTGATGGTCCTGTTCT<br>GCTGCCAGACAATCACTATCTGAGCACGCAAAGCGTTCTGTCTAAAGA<br>TCCGAACGAGAAACGC |

| | |
|---|---|
| | GATCATATGGTTCTGCTGGAGTTCGTAACCGCAGCGGGCATCACGCATG<br>GTATGGATGAACTGT ACAAATGATGA |
| *mScarlet* | ATGGTCAGTAAAGGAGAAGCTGTAATTAAAGAGTTTATGCGCTTCAAA<br>GTGCATATGGAAGGTTCCATGAACGGACATGAGTTCGAGATTGAAGGT<br>GAAGGTGAAGGACGCCCCTACGAGGGCACTCAAACTGCAAAGTTAAA<br>GGTTACAAAAGGAGGACCTCTGCCATTTTCGTGGGACATTCTGAGCCC<br>GCAGTTTATGTACGGCAGCCGCGCGTTCATCAAACATCCCGCTGACATC<br>CCAGACTATTACAAACAATCTTTCCCCGAAGGCTTTAAATGGGAACGC<br>GTGATGAACTTTGAAGATGGTGGCGCTGTGACTGTGACCCAGGACACT<br>TCATTAGAAGATGGAACCCTGATTTACAAGGTAAAGCTGCGCGGCACC<br>AACTTTCCCCCTGACGGACCTGTAATGCAGAAAAAAACAATGGGTTGG<br>GAGGCTAGTACAGAGCGTTTATACCCTGAGGACGGTGTCTTAAAAGGA<br>GACATCAAGATGGCGTTACGTCTTAAGGATGGTGGTCGCTATTTAGCTG<br>ACTTCAAGACCACTTATAAAGCAAAGAAGCCCGTCCAAATGCCTGGAG<br>CTTATAACGTTGACCGTAAGTTAGACATCACCTCACATAACGAGGATTA<br>CACAGTTGTCGAACAGTATGAGCGCTCAGAAGGCCGTCATTCGACTGG<br>TGGAATGGACGAACTGTATAAATAA |
| *mtrC* | ATGATGAACGCACAAAAATCAAAAATCGCACTGCTGCTCGCAGCAAGT<br>GCCGTCACAATGGCCTT<br>AACCGGCTGTGGTGGAAGCGATGGTAATAACGGCAATGATGGTAGTGA<br>TGGTGGTGAGCCAGCA<br>GGTAGCATCCAGACGTTAAACCTAGATATCACTAAAGTAAGCTATGAAA<br>ATGGTGCACCTATGGT<br>CACTGTTTTCGCCACTAACGAAGCCGACATGCCAGTGATTGGTCTCGC<br>AAATTTAGAAATCAAAA<br>AAGCACTGCAATTAATACCGGAAGGGGCGACAGGCCCAGGTAATAGCG<br>CTAACTGGCAAGGCTT<br>AGGCTCATCAAAGAGCTATGTCGATAATAAAAACGGTAGCTATACCTTT<br>AAATTCGACGCCTTCGA<br>TAGTAATAAGGTCTTTAATGCTCAATTAACGCAACGCTTTAACGTTGTTT<br>CTGCTGCGGGTAAATT<br>AGCAGACGGAACGACCGTTCCCGTTGCCGAAATGGTTGAAGATTTCGA<br>CGGCCAAGGTAATGCG<br>CCGCAATATACAAAAAATATCGTTAGCCACGAAGTATGTGCTTCTTGCC<br>ACGTAGAAGGTGAAAA<br>GATTTATCACCAAGCTACTGAAGTCGAAACTTGTATTTCTTGCCACACT<br>CAAGAGTTTGCGGATGG<br>TCGCGGCAAACCCCATGTCGCCTTTAGTCACTTAATTCACAATGTGCAT<br>AATGCCAACAAAGCTT |

| | |
|---|---|
| | GGGGCAAAGACAATAAAATCCCTACAGTTGCACAAAATATTGTCCAAG<br>ATAATTGCCAAGTTTGTC<br>ACGTTGAATCCGACATGCTCACCGAGGCAAAAAACTGGTCACGTATTC<br>CAACAATGGAAGTCTGT<br>TCTAGCTGTCACGTAGACATCGATTTTGCTGCGGGTAAAGGCCACTCTC<br>AACAACTCGATAACTC<br>CAACTGTATCGCCTGCCATAACAGCGACTGGACTGCTGAGTTACACAC<br>AGCCAAAACCACCGCA<br>ACTAAGAACTTGATTAATCAATACGGTATCGAGACTACCTCGACAATTA<br>ATACCGAAACTAAAGCA<br>GCCACAATTAGTGTTCAAGTTGTAGATGCGAACGGTACTGCTGTTGATC<br>TCAAGACCATCCTGCC<br>TAAAGTGCAACGCTTAGAGATCATCACCAACGTTGGTCCTAATAATGCA<br>ACCTTAGGTTATAGTGG<br>CAAAGATTCAATATTTGCAATCAAAAATGGAGCTCTTGATCCAAAAGCT<br>ACTATCAATGATGCTGG<br>CAAACTGGTTTATACCACTACTAAAGACCTCAAACTTGGCCAAAACGG<br>CGCAGACAGCGACACAGCATTTAGCTTTGTAGGTTGGTCAATGTGTTCT<br>AGCGAAGGTAAGTTTGTAGACTGTGCAGACCCTGCATTTGATGGTGTT<br>GATGTAACTAAGTATACCGGCATGAAAGCGGATTTAGCCTTTGCTACTT<br>TGTCAGGTAAAGCACCAAGTACTCGCCACGTTGATTCTGTTAACATGAC<br>AGCCTGTGCCAATTGCCACACTGCTGAGTTCGAAATTCACAAAGGCAA<br>ACAACATGCAGGCTTTGTGATGACAGAGCAACTATCACACACCCAAGA<br>TGCTAACGGTAAAGCGATTGTAGGCCTTGACGCATGTGTGACTTGTCAT<br>ACTCCTGATGGCACCTATAGCTTTGCCAACCGTGGTGCGCTAGAGCTAA<br>AACTACACAAAAAACACGTTGAAGATGCCTACGGCCTCATTGGTGGCA<br>ATTGTGCCTCTTGTCACTCAGACTTCAACCTTGAGT<br>CTTTCAAGAAGAAAGGCGCATTGAATACTGCCGCTGCAGCAGATAAAA<br>CAGGTCTATATTCTACG<br>CCGATCACTGCAACTTGTACTACCTGTCACACAGTTGGCAGCCAGTAC<br>ATGGTCCATACGAAAGA<br>AACCCTGGAGTCTTTCGGTGCAGTTGTTGATGGCACAAAAGATGATGC<br>TACCAGTGCGGCACAG<br>TCAGAAACCTGTTTCTACTGCCATACCCCAACAGTTGCAGATCACACTA<br>AAGTGAAAATGTAA |
| *mtrA* | ATGAAGAACTGCCTAAAAATGAAAAACCTACTGCCGGCACTTACCATC<br>ACAATGGCAATGTCTGC<br>AGTTATGGCATTAGTCGTCACACCAAACGCTTATGCGTCGAAGTGGGAT<br>GAGAAAATGACGCCAG |

| | |
|---|---|
| | AGCAAGTCGAAGCCACCTTAGATAAGAAGTTTGCCGAAGGCAACTACT CCCCTAAAGGCGCCGA TTCTTGCTTGATGTGCCATAAGAAATCCGAAAAAGTCATGGACCTTTTC AAAGGTGTCCACGGTG CGATTGACTCCTCTAAGAGTCCAATGGCTGGCCTGCAATGTGAGGCAT GCCACGGCCCACTGGG TCAGCACAACAAAGGCGGCAACGAGCCGATGATCACTTTTGGTAAGCA ATCAACCTTAAGTGCCGACAAGCAAAACAGCGTATGTATGAGCTGTCA CCAAGACGATAAGCGTATGTCTTGGAATGGCGGTCACCATGACAATGC CGATGTTGCTTGTGCTTCTTGTCACCAAGTACACGTCGCAAAAGATCCT GTGTTATCTAAAAACACGGAAATGGAAGTCTGTACTAGCTGCCATACAA AGCAAAAAGCGGATATGAATAAACGCTCAAGTCACCCACTCAAATGGG CACAAATGACCTGTAGCGACTGTCACAATCCCCAT GGGAGCATGACAGATTCCGATCTTAACAAGCCTAGCGTGAATGATACCT GTTATTCCTGTCACGC CGAAAAACGCGGCCCAAAACTTTGGGAGCATGCACCCGTCACTGAGA ATTGTGTCACTTGCCAC AATCCTCACGGTAGTGTGAATGACGGTATGCTGAAAACCCGTGCGCCA CAGCTATGTCAGCAAT GTCACGCCAGCGATGGCCACGCCAGCAACGCCTACTTAGGTAACACTG GATTAGGTTCAAATGT CGGTGACAATGCCTTTACTGGTGGAAGAAGCTGCTTAAATTGCCATAGT CAGGTTCATGGTTCTA ACCATCCATCTGGCAAGCTATTACAGCGCTAA |
| *mtrB* | ATGAAATTTAAACTCAATTTGATCACTCTAGCGTTATTAGCCAACACAG GCTTGGCCGTCGCTGCTGATGGTTATGGTCTAGCGAATGCCAATACTGA AAAAGTGAAATTATCCGCATGGAGCTGTAAAGGCTGCGTCGTTGAAAC GGGCACATCAGGCACTGTGGGTGTCGGTGTCGGTTATAACAGCGAAGA GGATATTCGCTCTGCCAATGCCTTTGGTACATCCAATGAAGTGGCGGGT AAATTTGATGCCGATTTAAACTTTAAAGGTGAAAAGGGTTATCGTGCCA GTGTTGATGCTTATCAACTCGGTATGGATGGCGGTCGCTTAGATGTCAA TGCGGGCAAACAAGGCCAGTACAACGTCAATGTGAACTATCGCCAAAT TGCTACCTACGACAGCAATAGCGCCCTATCGCCCTACGCGGGTATTGGT GGCAATAACCTCACGTTACCGGATAACTGGATAACAGCAGGTTCAAGC AACCAAATGCCACTCTTGATGGACAGCCTCAATGCCCTCGAACTCTCA CTTAAACGTGAGCGCACGGGGTTGGGATTTGAATATCAAGGTGAATCC CTGTGGAGCACCTATGTTAACTACATGCGTGAAGAGAAAACCGGCTTA AAACAAGCCTCTGGTAGCTTCTTCAACCAATCGATGATGTTAGCAGAG CCGGTGGATTACACCACTGACACCATTGAAGCGGGTGTCAAACTCAAG GGTGATCGTTGGTTTACCGCACTCAGTTACAATGGGTCAATATTCAAAA |

ACGAATACAACCAATTGGACTTTGAAAATGCTTTTAACCCCACCTTTGG
TGCTCAAACCCAAGGTACGATGGCACTCGATCCGGATAACCAGTCACA
CACCGTGTCGCTGATGGGACAGTACAACGATGGCAGCAACGCACTGTC
GGGTCGTATTCTGACCGGACAAATGAGCCAAGATCAGGCGTTAGTGAC
GGATAACTACCGTTATGCTAATCAGCTCAATACCGATGCCGTCGATGCC
AAAGTCGATCTACTGGGTATGAACCTGAAAGTCGTTAGCAAAGTGAGC
AATGATCTTCGCTTAACAGGTAGTTACGATTATTACGACCGTGACAATA
ATACCCAAGTAGAAGAATGGACTCAGATCAGCATCAACAATGTCAACG
GTAAGGTGGCTTATAACACCCCTTACGATAATCGTACGCAACGCTTTAA
AGTTGCCGCAGATTATCGCATTACCCGCGATATCAAACTCGATGGTGGT
TATGACTTCAAACGTGACCAACGTGATTATCAAGACCGTGAAACCACG
GATGAAAATACCGTTTGGGCCCGTTTACGTGTAAACAGCTTCGATACTT
GGGACATGTGGGTAAAAGGCAGTTACGGTAACCGTGACGGCTCACAAT
ACCAAGCGTCTGAATGGACCTCTTCTGAAACCAACAGCCTGTTACGTA
AGTACAATCTGGCTGACCGTGACAGAACTCAAGTCGAAGCACGGATCA
CCCATTCGCCATTAGAAAGCCTGACTATCGATGTTGGTGCCCGTTACGC
GTTAGATGATTATACCGATACTGTGATTGGATTAACTGAGTCAAAAGAC
ACCAGTTATGATGCCAACATCAGTTATATGATCACCGCTGACTTACTGGC
AACCGCCTTCTACAATTACCAAACCATTGAGTCTGAACAGGCGGGTAG
CAGCAATTACAGCACCCCAACGTGGACAGGCTTTATAGAAGATCAGGT
AGATGTGGTCGGTGCAGGTATCAGCTACAACAATCTGCTGGAGAACAA
GTTACGCCTAGGACTGGACTACACCTATTCCAACTCCGACAGTAACACT
CAAGTCAGACAAGGTATCACTGGCGACTATGGTGATTATTTTGCCAAAG
TGCATAACATTAACTTATACGCTCAATATCAAGCCACCGAGAAACTCGC
GCTGCGCTTCGATTACAAAATTGAGAACTATAAGGACAATGACGCCGC
AAATGATATCGCCGTTGATGGCATTTGGAACGTCGTAGGTTTTGGTAGT
AACAGCCATGACTACACCGCACAAATGCTGATGCTGAGCATGAGTTAC
AAACTCTAA

**Table S3:** Nonlinear (sigmoidal) gene expression fit parameters for products of *sfgfp* expression (fluorescence) or EET gene expression (storage modulus).
Parameters were fit using an activating or deactivating 4-parameter sigmoidal model (Hill Function) in Prism 9.

| Plasmid Name | Measured Outcome | n | $k_{1/2}$ (nM) | $y_{max}$ (units) | $y_{min}$ (units) | Dynamic Range | Goodness of Fit ($R^2$) |
|---|---|---|---|---|---|---|---|
| pAT3 | Fluorescence | 2.125 | 3.073 | 1.479 (REU) | 0.01799 (REU) | 1.461 | .98 |
| pAT4 | Fitted Rate Constant (Ferrozine) | 1.348 | 3.160 | 1.647 ($h^{-1}$) | .2480 ($h^{-1}$) | 1.399 | .90 |
| pAT4 | B2 Value (CuAAC Rate) | .8301 | 4.550 | 135.7 (fluorescence a.u./$h^2$) | 63.24 (fluorescence a.u./$h^2$) | 72.44 | .52 |
| pAT4 | Storage Modulus (1% ACAG, 5% PEGMA) | 1.236 | 3.917 | 1441 (Pa) | 4028 (Pa) | 2586 | .91 |
| pAT4 | Storage Modulus (5% PEGMA) | 2.586 | 2.065 | 133.1 (Pa) | 1.070 (Pa) | 132 | .75 |
| pAT3NOT | Fluorescence | -1.843 | .4624 | 77,602 (Fluor./OD) | 6,404 (Fluor./OD) | 71,198 | .99 |
| PAT4NOT | Fitted Rate Constant (Ferrozine) | -1.004 | 5.458 | .923 ($h^{-1}$) | .1004 ($h^{-1}$) | .8019 | .99 |
| pAT3mScarlet | Fluorescence | .5649 | 2.715 | 276,260 (Fluor./OD) | -7338 (Fluor./OD) | 283,597 | .93 |

**Supplementary Figures**

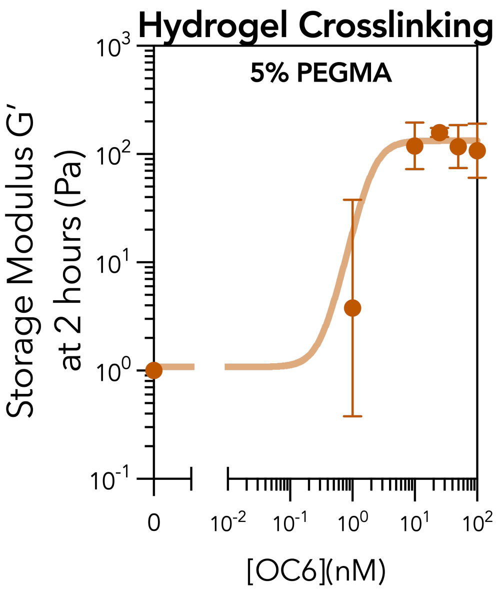


**Figure S1.** EET-driven crosslinking of 5% methacrylated PEG hydrogels. Storage moduli of 5 wt% methacrylated PEG and 1 wt% acrylated agarose co- polymer networks cross-linked via variably induced S.oneidensis ΔMtr containing pAT4 driving atom-transfer radical polymerization (ATRP). Moduli were measured using shear rheometry after swelling. Individual replicates were fit to a four-parameter activating Hill function with weighted error. All data represent the mean ± SEM of the appropriate measurement for n = 3 biological replicates.

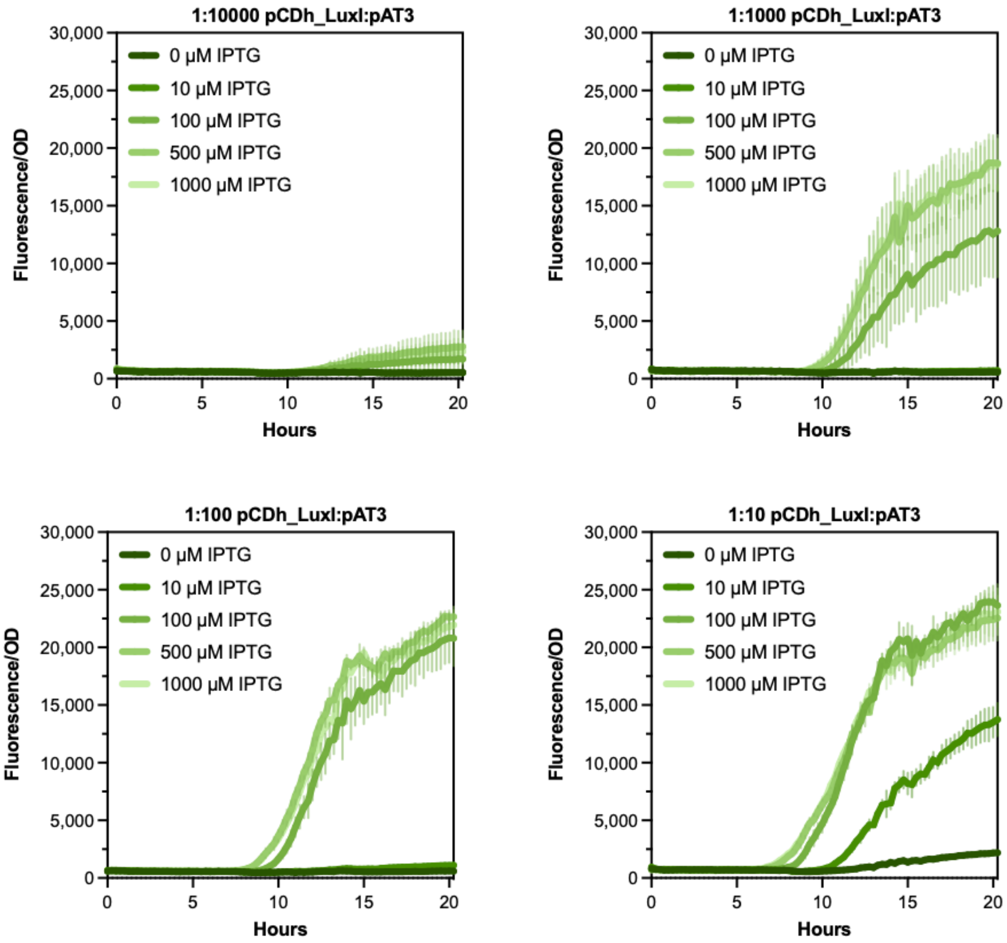


**Figure S2.** Raw pCDh_LuxI + pAT3 co-culture fluorescence kinetics.

Variable sender cell induction in co-culture at variable ratios with sfgfp receiver cells. Fluorescence and $OD_{600}$ were measured every 15 minutes over 20 hours of aerobic growth at 30°C. Fluorescence values were OD-normalized at each time point. Endpoint fluorescence was used to create the heatmap in Figure 3.2.b. Darker shades of green correspond to higher levels of sender cell induction with IPTG. Cultures with a higher level of sender cell induction and/or a higher sender:receiver ratio demonstrate a higher endpoint fluorescence value as well as faster reaction kinetics. After 20 hours, even samples without induction at the smallest ratio of 1:10,000 show some sfgfp expression likely due to the slight leakiness of the *Ptac* promoter controlling LuxI expression in the sender cells. n=3 for each ratio and inducer condition.

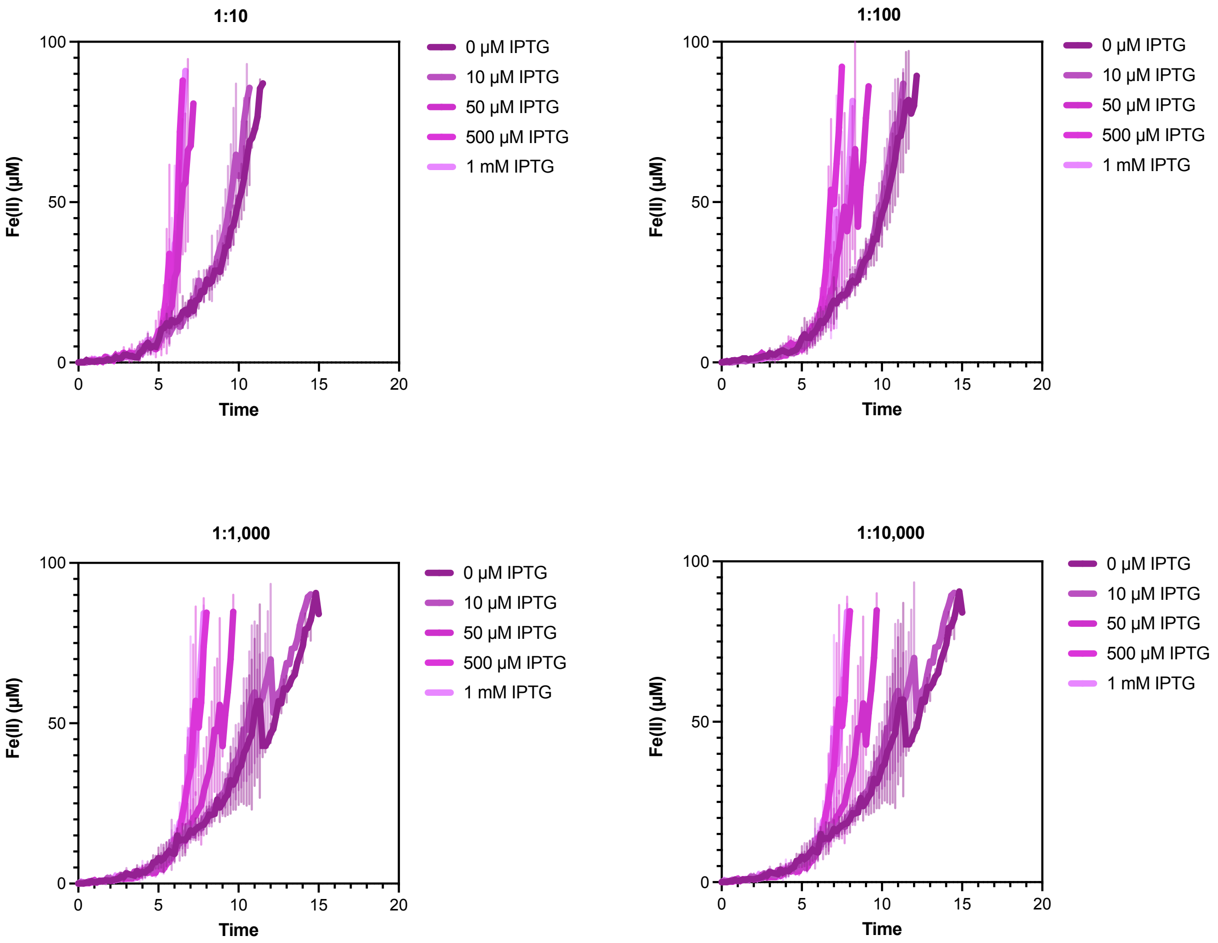


**Figure S3.** Raw pCDh_LuxI + pAT4 co-culture Fe(III) reduction kinetics.

Variable sender cell induction in co-culture at variable ratios with EET receiver cells. Absorbance at 562 nm was measured every 15 minutes over 14 hours during anaerobic growth at 30°C. Fluorescence values were OD-normalized at each time point. In situ Fe(III) reduction data were fit to a Monod-type model as described in the Methods to obtain fitted rate which were used to create the heatmap in Figure 3.2.c. Darker shades of purple correspond to higher levels of sender cell induction with IPTG. Sender cells were in a **Δ**Mtr background. Cultures with a higher level of sender cell induction and/or a higher sender:receiver ratio demonstrate higher rates of EET, with an almost 10 hour difference in the time needed to saturate the absorbance measurement. After 20 hours, even samples without induction at the smallest ratio of 1:10,000 show Fe(III) expression likely due to the slight leakiness of the *Ptac* promoter controlling LuxI expression in the sender cells and background EET processes that don't rely on the Mtr pathway. n=3 for each ratio and inducer condition.

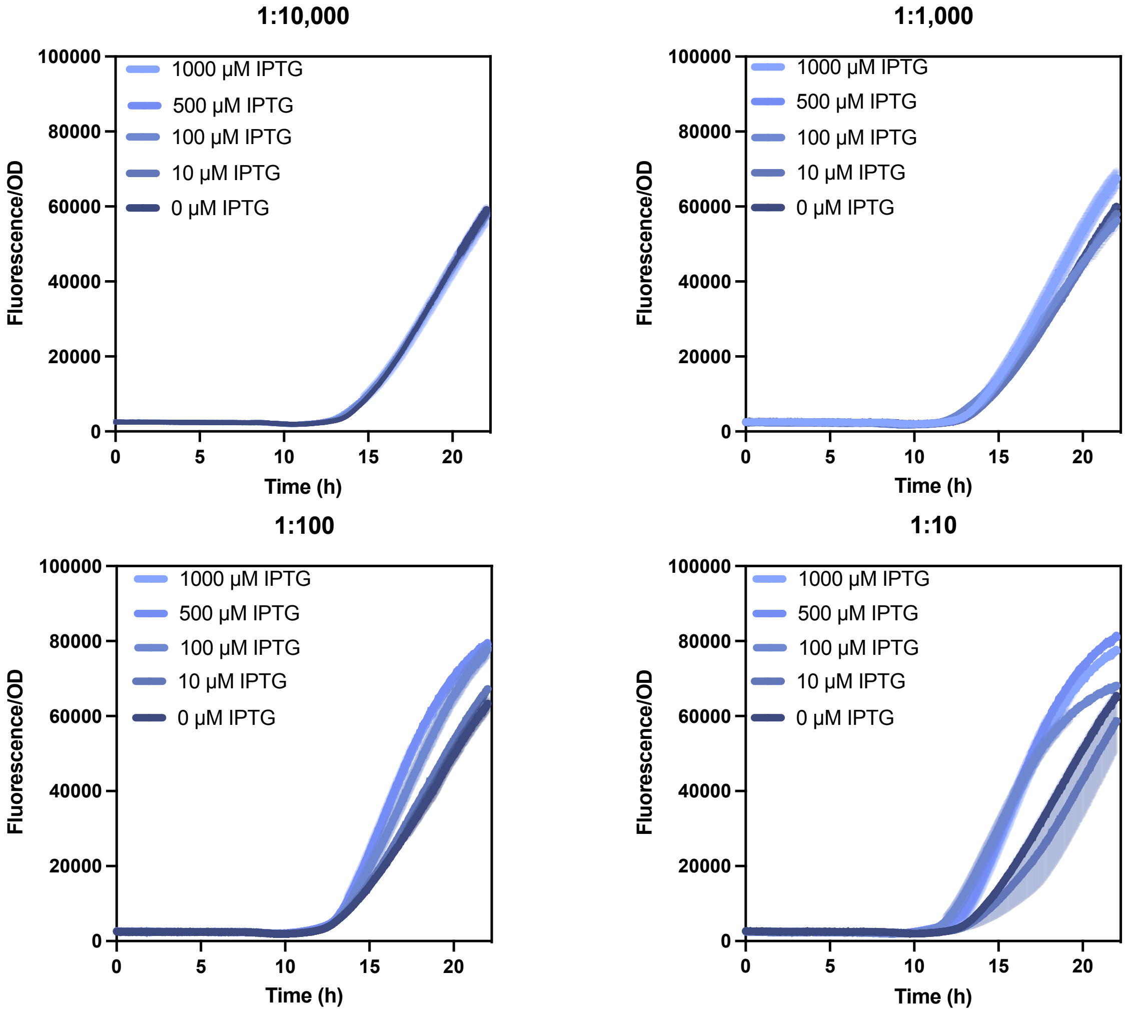


**Figure S4.** Raw pCDh_LuxI + pAT4 co-culture CuAAC probe reaction kinetics.

Variable sender cell induction in co-culture at variable ratios with EET receiver cells. Fluorescence was measured every 15 minutes over 20 hours during anaerobic growth at 30°C to track the CuAAC catalyzed formation of a fluorescent probe. Fluorescence values were OD-normalized at each time point. Fluorescence data were fit to a second-order polynomial function as explainnend in the Methods to obtain the fitted rates (B2 value) used to create the heatmap in Figure 3.2.d. Darker shades of blue correspond to higher levels of sender cell induction with IPTG. Sender cells were in a **Δ**Mtr background. Cultures with a higher level of sender cell induction and/or a higher sender:receiver ratio demonstrate higher rates and yield of CuAAC product formation. After 20 hours, even samples without induction at the smallest ratio of 1:10,000 show conversion likely due to the slight leakiness of the *Ptac* promoter controlling LuxI expression in the sender cells and background EET processes that don't rely on the Mtr pathway. n=3 for each ratio and inducer condition.

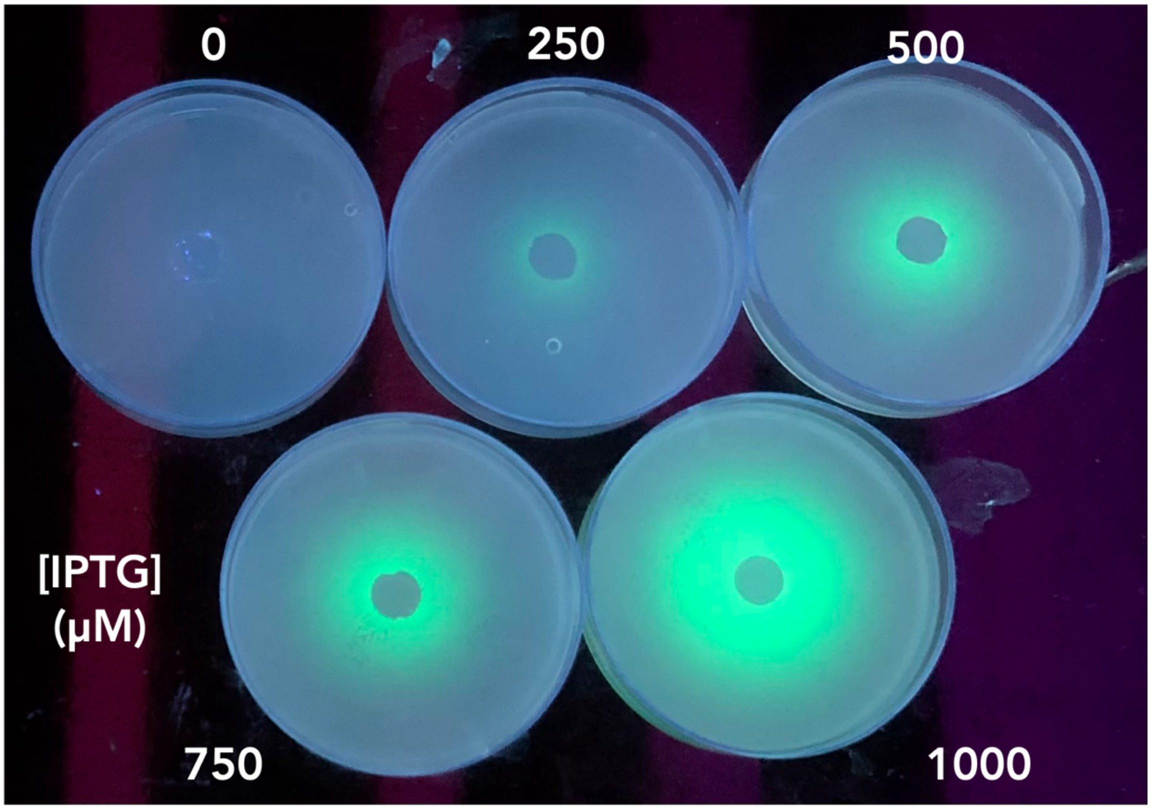


**Figure S5.** Top-down view of pAT3 fluorescence patterns with variable sender cell induction. SBM plates supplemented with cas amino acids, Wolfe's mineral mix, lactate, fumarate, and kanamycin were used as a platform for a .7% agarose lawn with the same media composition containing pAT3 receiver cells. Following 18-hour aerobic incubation at 30°C with sender discs containing concentrated sender cells, the plates were photographed on an UV transilluminator. The IPTG concentration used to induce the sender cells is displayed adjacent to each plate.

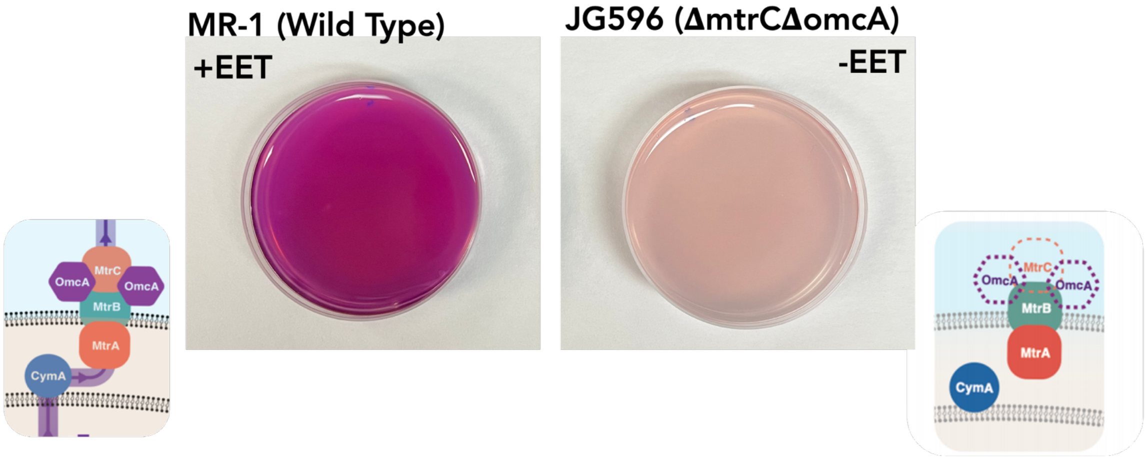


**Figure S6:** Top-down view of wild-type and ***Δ****mtrC S.oneidensis* ferrozine reduction patterns. SBM plates supplemented with cas amino acids, lactate, and fumarate were used as a platform for a .7% agarose lawn with the same media composition containing either wild-type or EET-deficient strains. Following 18-hour aerobic incubation at 30°C, 3 mL of .7% SBM agar containing 2 mM Fe(III) citrate and 1 mM ferrozine was poured on the plates. The plates incubated at room temperature for 10 minutes before being imaged. The illustrations on either side depict the EET machinery of each sample. As expected, the wild-type showed a quick and significant purple color shift, while the knockout took significantly longer to reach the same intensity. These data serve as a baseline for the expected behavior of patterns of EET expression in the presence of iron and ferrozine.

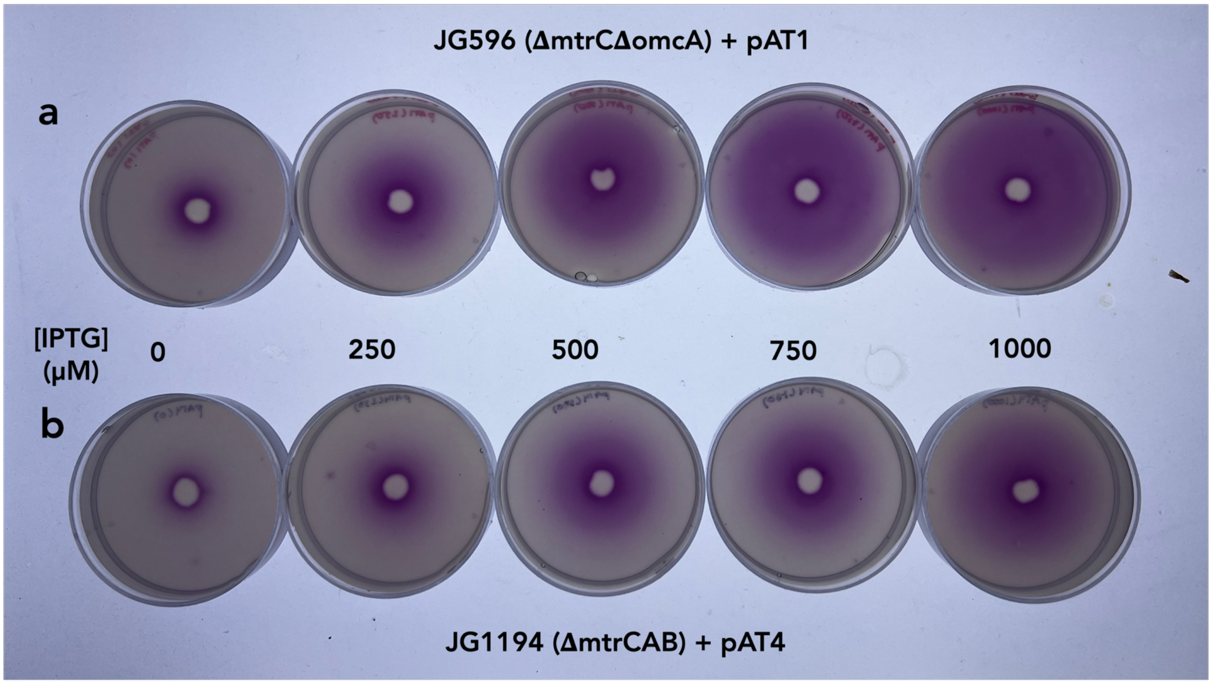


**Figure S7.** Top-down view of pAT1 and pAT4 Fe(III) reduction patterns with variable sender cell induction.

SBM plates supplemented with cas amino acids, lactate, fumarate, and kanamycin were used as a platform for a .7% agarose lawn with the same media composition containing **a.** pAT1 or **b.** pAT4 receiver cells. Following 18-hour aerobic incubation at 30°C with sender discs containing concentrated sender cells, 3 mL of .7% SBM agar containing 2 mM Fe(III) citrate and 1 mM ferrozine was poured on the plates. The plates incubated at room temperature for 10 minutes before being imaged on a light table. The IPTG concentration used to induce the sender cells is displayed adjacent to each plate. The pAT4 cells show slower and smaller levels of response to the same OC6 levels compared to pAT1. pAT4 allows for less background reduction since the MtrCAB pathway is genomically knocked out, and expression of the polycistronic *mtrCAB* operon on the plasmid is more metabolically burdensome.

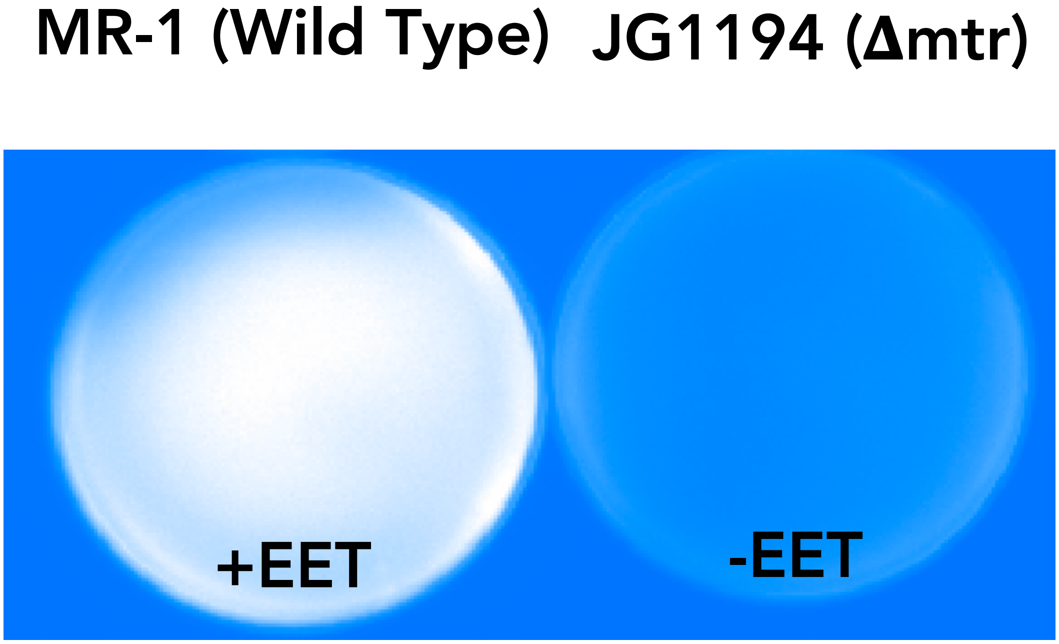


**Figure S8:** Top-down view of wild-type and Δ*mtrCAB* CuAAC probe conversion patterns. SBM plates supplemented with cas amino acids, lactate, and fumarate were used as a platform for a .7% agarose lawn with the same media composition containing either wild-type or EET-deficient strains. Following 18-hour aerobic incubation at 30°C, 3 mL of .7% SBM agar containing CuAAC reaction reagents was poured on the plates. The plates incubated anaerobically at 30°C for 5 hours before being imaged on a gel imager. As expected, the wild-type showed a significant fluorescence signal indicative of successful CuAAC catalysis. The knockout shows virtually no fluoresennce. These data serve as a baseline for the expected behavior of patterns of EET expression in the presence of CuAAC reagents.

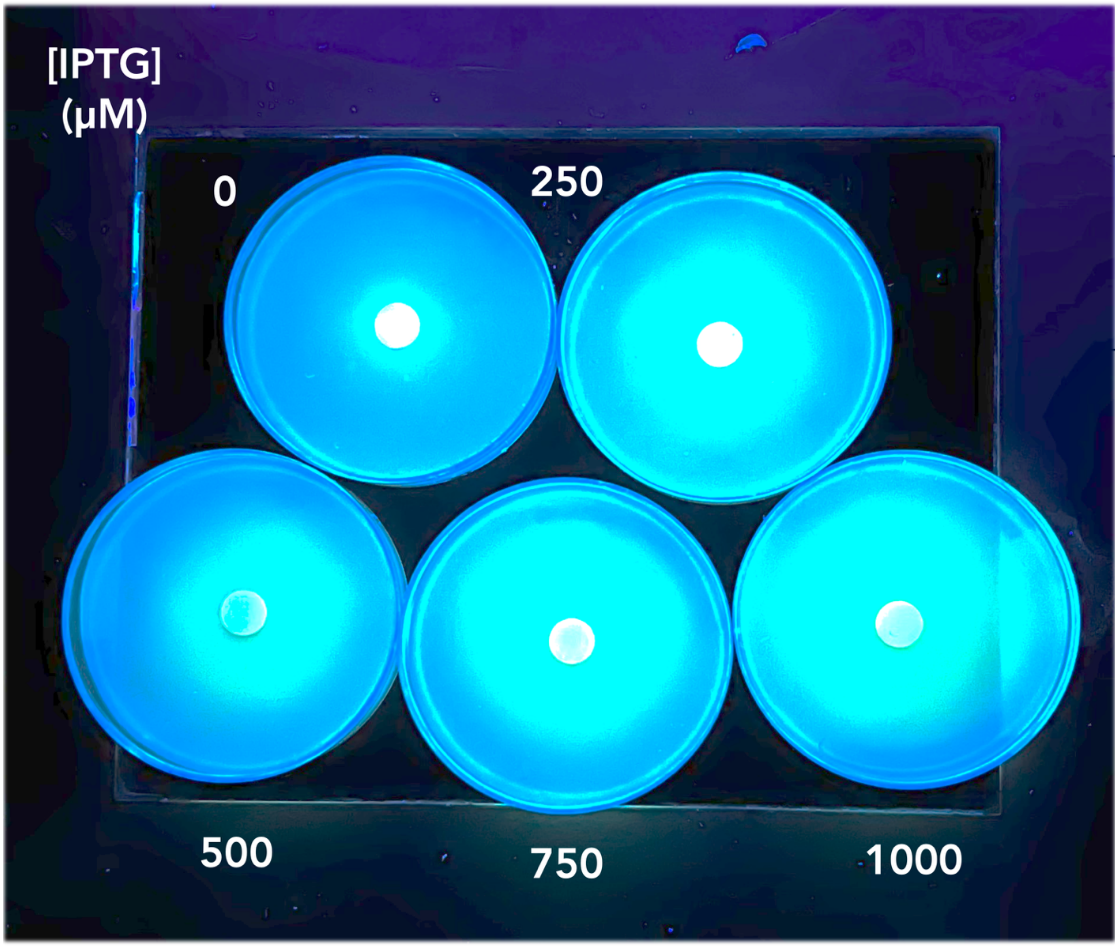


**Figure S9.** Top-down view of pAT4 CuAAC probe biocatalysis with variable sender cell induction.

SBM plates supplemented with cas amino acids, lactate, fumarate, and kanamycin were used as a platform for a .7% agarose lawn with the same media composition containing **a.** pAT1 or **b.** pAT4 receiver cells. Following 18-hour aerobic incubation at 30°C, 3 mL of .7% SBM agar containing CuAAC reaction reagents was poured on the plates. The plates incubated anaerobically at 30°C for 5 hours before being imaged on a UV transilluminator. The IPTG concentration used to induce the sender cells is displayed adjacent to each plate.

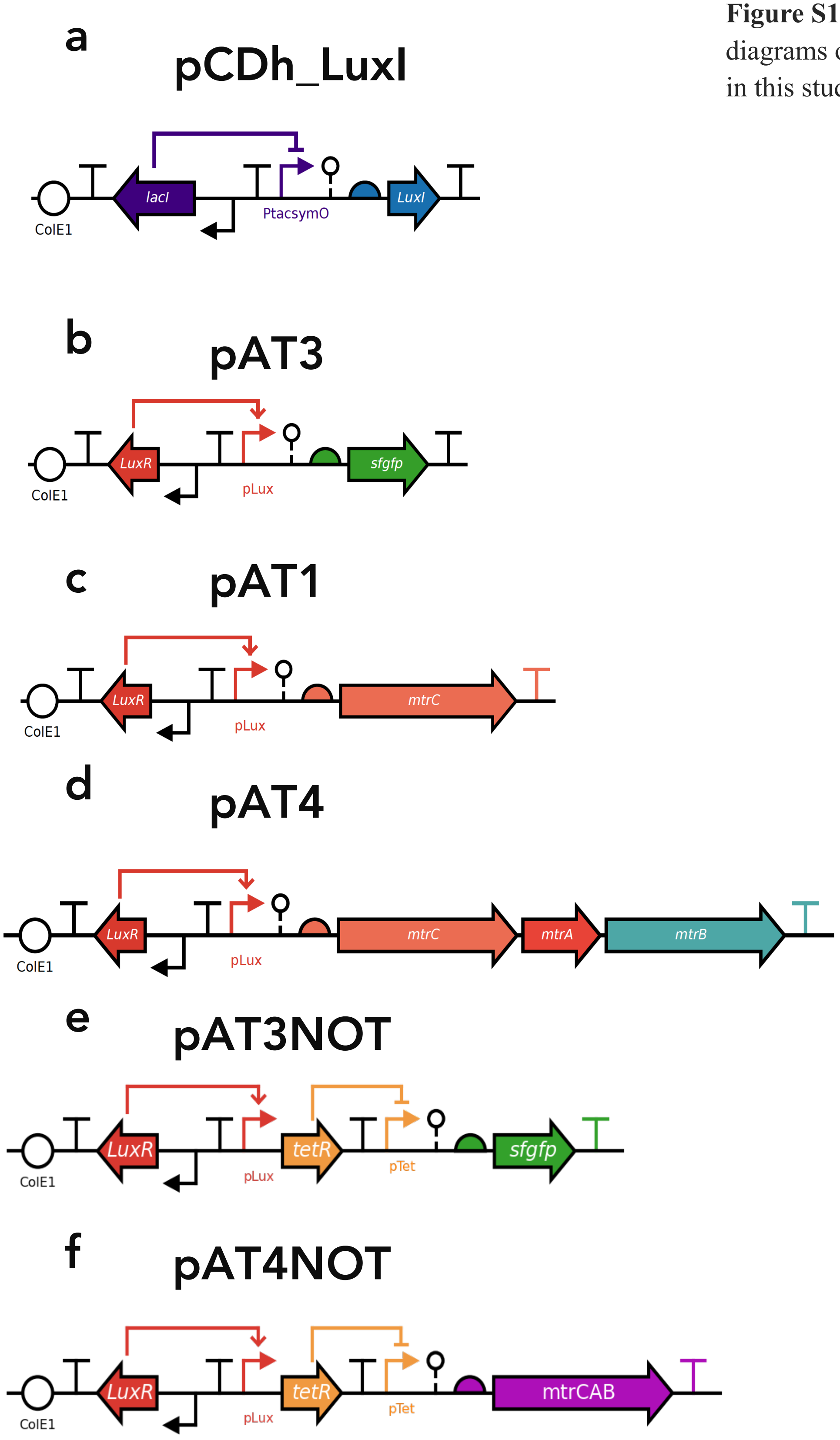


**Figure S10.** Gene circuit diagrams of the plasmids used in this study

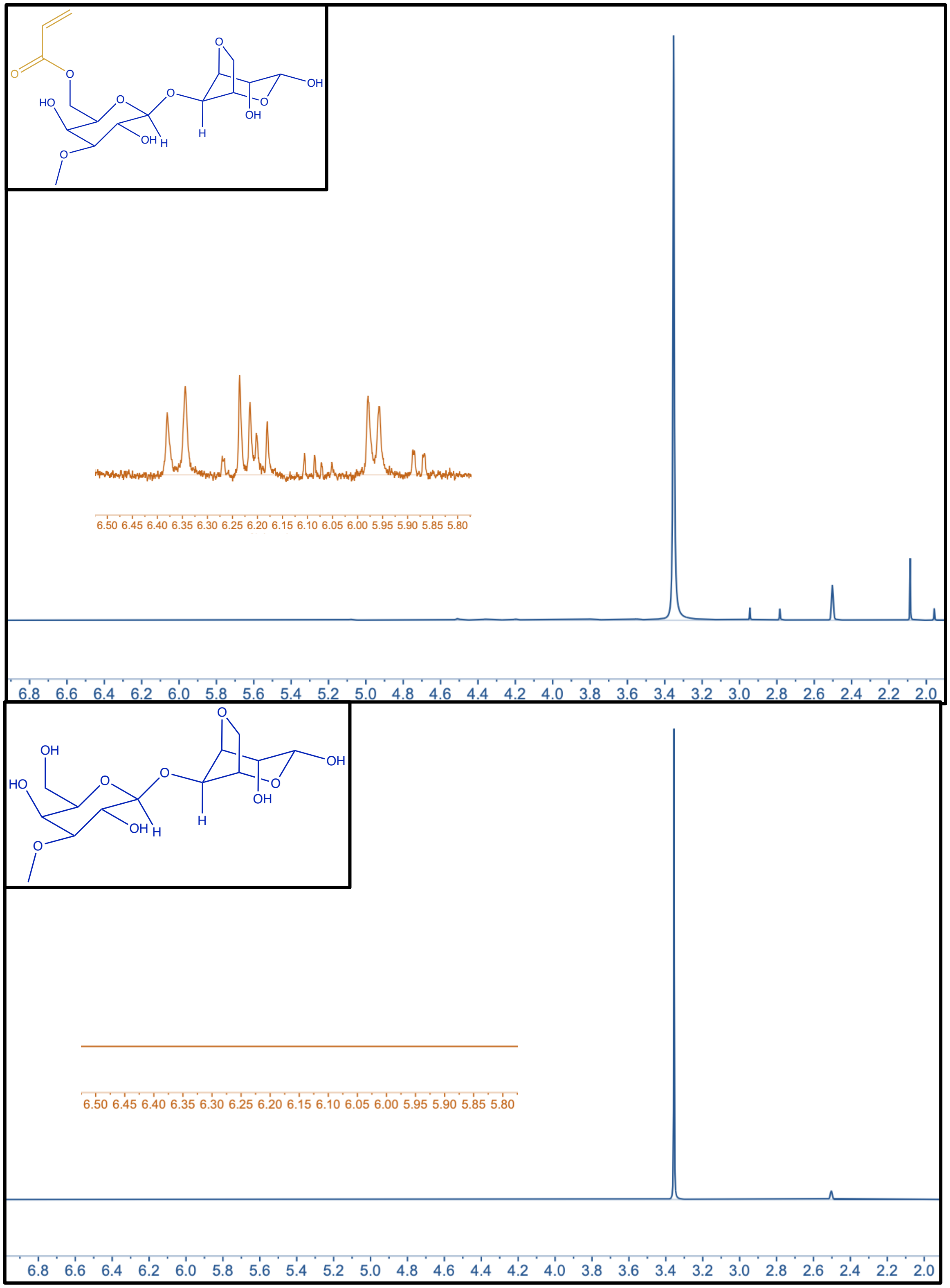


**Figure S11:** $^{1}$H-NMR spectra of acrylated (top) and unmodified (bottom) agarose.

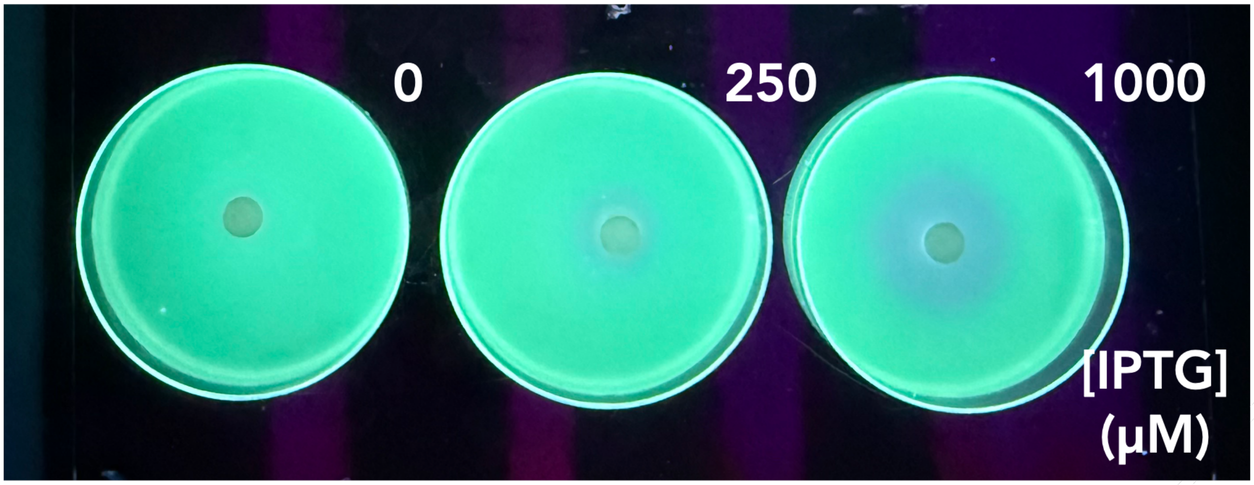


**Figure S12.** Top-down view of pAT3NOT fluorescence patterns with variable sender cell induction.

SBM plates supplemented with cas amino acids, Wolfe's mineral mix, lactate, fumarate, and kanamycin were used as a platform for a .7% agarose lawn with the same media composition containing pAT3 receiver cells. Following 18-hour aerobic incubation at 30°C with sender discs containing concentrated sender cells, the plates were photographed on an UV transilluminator. The IPTG concentration used to induce the sender cells is displayed adjacent to each plate.

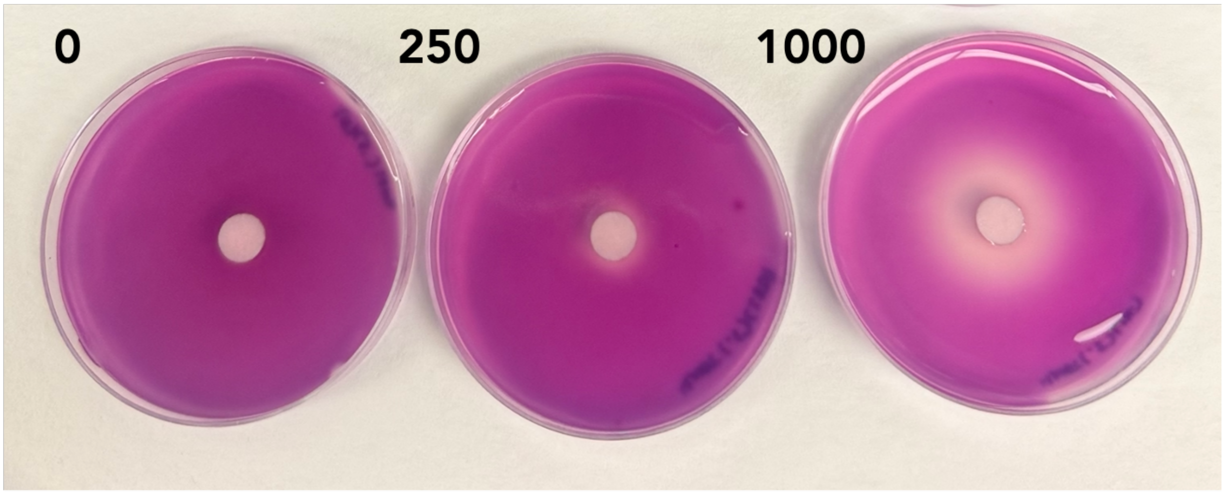


**Figure S13.** Top-down view of pAT4NOT Fe(III) reduction patterns with variable sender cell induction.

SBM plates supplemented with cas amino acids, lactate, fumarate, and kanamycin were used as a platform for a .7% agarose lawn with the same media composition containing pAT4 receiver cells. Following 18-hour aerobic incubation at 30°C with sender discs containing concentrated sender cells, 3 mL of .7% SBM agar containing 2 mM Fe(III) citrate and 1 mM ferrozine was poured on the plates. The plates incubated at room temperature for 10 minutes before being imaged on a light table. The IPTG concentration used to induce the sender cells is displayed adjacent to each plate.

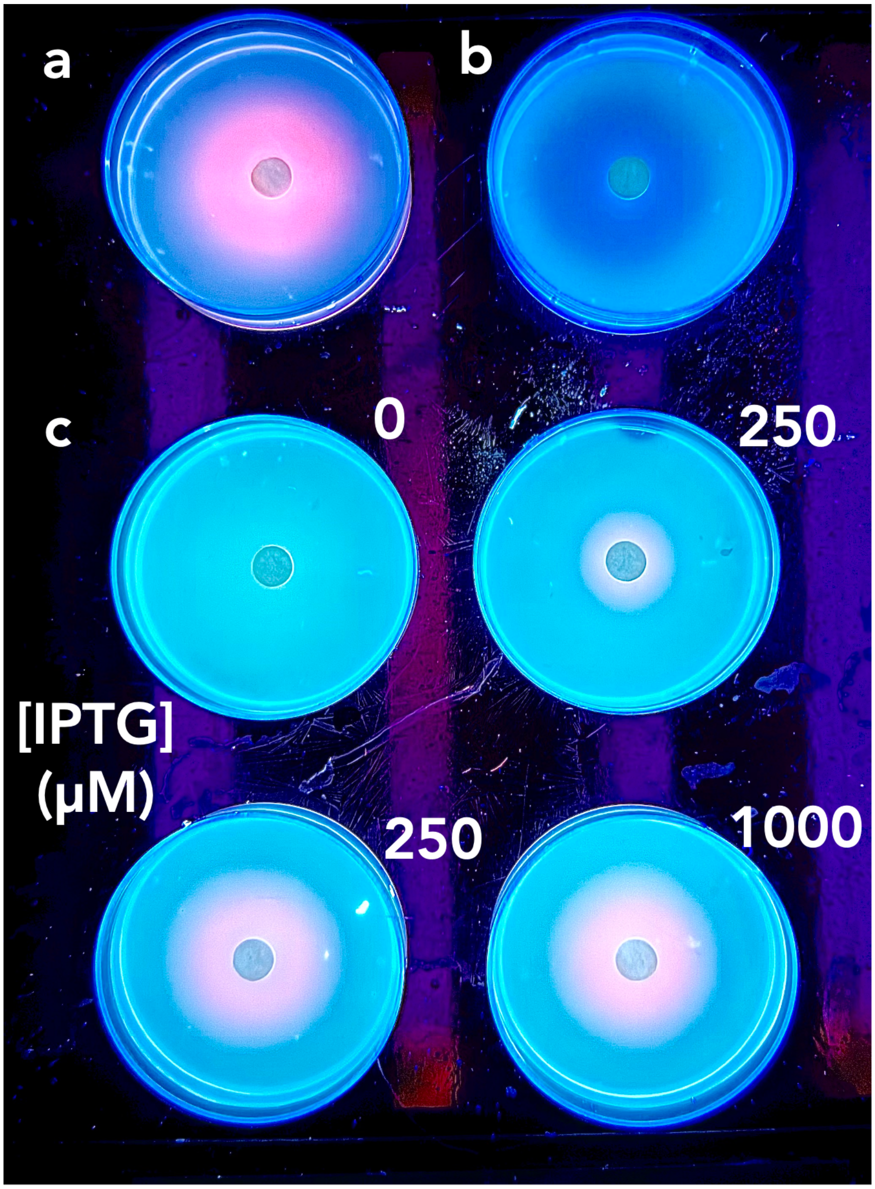


**Figure S14:** Top-down view of pAT3mScarlet and pAT3NOT fluorescence patterns with variable sender cell induction.

SBM plates supplemented with cas amino acids, Wolfe's mineral mix, lactate, fumarate, and kanamycin were used as a platform for a .7% agarose lawn with the same media composition containing a) pAT3mScarlet, b) pAT3NOT, or c) 1:1 pAT3mScarlet:pAT3NOT receiver cells. Following 18-hour aerobic incubation at 30°C with sender discs containing concentrated sender cells, the plates were photographed on an UV transilluminator. The IPTG concentration used to induce the sender cells is displayed adjacent to each plate.
